# Age-related decline in neural phase-locking to envelope and temporal fine structure revealed by frequency following responses: A potential signature of cochlear synaptopathy impairing speech intelligibility

**DOI:** 10.1101/2024.12.11.628010

**Authors:** Emmanuel Ponsot, Pauline Devolder, Ingeborg Dhooge, Sarah Verhulst

## Abstract

Assessing the contribution of cochlear synaptopathy (CS) to the variability in speech-in-noise intelligibility among individuals remains a challenge. While several studies have proposed biomarkers for CS based on neural phase-locking to the temporal envelope (ENV), fewer have investigated how CS affects the coding of temporal fine structure (TFS), despite its crucial role in speech-in-noise perception. In this study, we specifically examined whether TFS-based markers of CS could be derived from electrophysiological responses and psychophysical detection thresholds of spectral modulation (SM) in a complex tone, which serves as a parametric model of speech. We employed an integrated approach, combining psychophysical testing with frequency-following response (FFR) measurements in three groups of participants: young normal-hearing (yNH), older normal-hearing (oNH), and older hearing-impaired (oHI) individuals. We expanded on previous work by assessing phase-locking to both ENV, using a 4 kHz rectangular amplitude-modulated (RAM) tone, and TFS, using a low-frequency (<1.5 kHz) SM complex tone. Overall, FFR results showed significant reductions in neural phase-locking to both ENV and TFS components with age and hearing loss. Specifically, the strength of TFS-related FFRs, particularly the component corresponding to the harmonic closest to the peak of the spectral envelope (∼500 Hz), was negatively correlated with age, even after adjusting for audiometric thresholds. This TFS marker also correlated with ENV-related FFRs derived from the RAM tone, suggesting a shared decline in phase-locking capacity across low and high cochlear frequencies. Computational simulations of the auditory periphery indicated that the observed FFR strength reduction with age is consistent with approximately 50% loss of auditory nerve fibers, aligning with histopathological data. However, the TFS-based FFR marker did not account for variability in speech intelligibility observed in the same participants. Psychophysical measurements showed no age-related effects and were unrelated to the TFS-based FFR marker, highlighting the need for further psychophysical research to establish a behavioral counterpart. Altogether, our results demonstrate that FFRs to vowel-like stimuli can serve as a complementary electrophysiological marker for assessing neural coding fidelity to stimulus TFS. This approach could provide a valuable tool for better understanding the impact of CS on an important coding dimension for speech-in-noise perception.

## Introduction

Understanding speech in noisy environments is at the core of human social interactions; yet, many individuals present significant difficulties in these situations. Speech-in-noise (SPiN) performance is shown to be impaired by multiple factors which degrade sensory or cognitive processes, from audiometric thresholds to selective attention capacities (e.g., Parbery-Clark et al., 2009; Oberfeld & Kloeckner-Nowotny, 2016; Varnet et al., 2021), but recent studies demonstrated that these factors only provide a limited account to understand listeners’ performance variability (Ruggles et al., 2011; Bharadwaj et al., 2015). A striking illustration is provided by the large-scale study conducted by Varnet et al. (2021) wherein over 500 individuals of various ages and audiometric thresholds performed a very simplified speech-in-noise perception task (monaural, simple sentences embedded in speech-shaped noise). Listeners with comparable audiometric deficits exhibited a huge variability of speech-in-noise performance, even among listeners with clinically-normal hearing thresholds, despite the simplicity of adopted speech material which was far from the complexity of real cocktail-party-like listening and thus minimizing the contribution of linguistic and cognitive factors. Additionally, a recent study based on a database of ∼100k individuals who consulted an audiology department for various hearing-in-noise problems led to the estimate that more than 10% had clinically normal audiograms (Parthasarathy et al., 2020). Recent research studies are hence focusing on the role of sensory coding deficits different from hearing sensitivity for explaining this variability.

There are two research paths that have been followed over the past decades to unravel the origins of individual differences in speech intelligibility deficits. The first track focuses on supra-threshold psychoacoustic tasks that capture the individual sensitivity to detecting spectro-temporal sound modulations (STM; Bernstein et al., 2013, 2016; Mehraei et al, 2014, Miller et al., 2018). STMs constitute first-order models of speech units that are vowels and formant transitions (see Fig. 1) critical for speech intelligibility (Elliott and Theunissen, 2009). Several studies investigating STM detection in hearing-impaired (HI) listeners have reported correlations between STM detection thresholds (smallest detectable modulation depth) and SPiN scores, even after accounting for differences in audiometric thresholds (Bernstein et al., 2013, 2016; Mehraei et al, 2014, Miller et al., 2018). Because these correlations were specifically driven by STM signals with low-frequency carriers (<1 kHz) and observed with spectrally-modulated (SM) signals, i.e. that do not carry temporal modulation, it was hypothesized that the variability in SPiN performances reflects an impaired ability to use TFS information in the stimulus waveform (Miller et al., 2018). From an audiological point of view, STM stimuli became appealing candidates for providing a better characterization of auditory deficits compared to pure-tone audiometry; clinical tests and large-scale studies are therefore currently conducted in that direction (Zaar et al., 2023a, 2023b, 2023c). Interestingly, studies investigating STM perception in normal-hearing (NH) listeners have reported large interindividual differences in detection thresholds (Oetjen and Verhey, 2015, 2017), but these studies did not address the question of whether this variability would relate to TFS-processing deficits and further, to potential differences in SPiN scores. However, while psychoacoustic S(T)M sensitivity may be a promising predictor of SPiN, it cannot directly be used to quantify or treat the pathophysiology of SPiN deficits.

**Figure 1.**
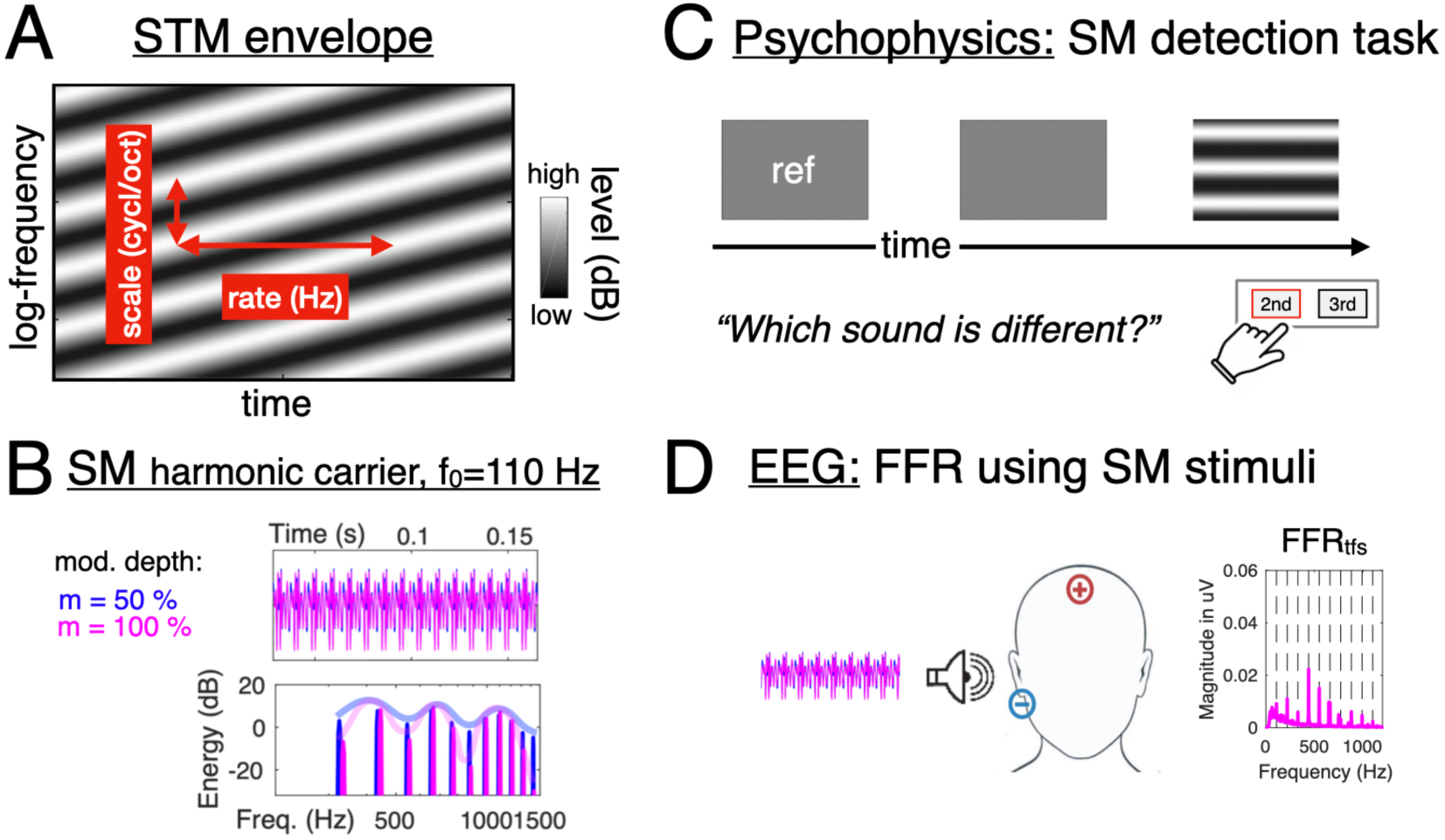
(A) Spectrogram representation of a STM stimulus used in the present study, showing the rate and scale parameters. (B) SM stimuli with harmonic carrier (SMharm) used in the psychophysical detection tasks as well as for the FFR measurements illustrated in (C) and (D), respectively.

Therefore, a second line of research focused on the role of cochlear synaptopathy (CS) for speech intelligibility. CS refers to the loss or damage of synapses between inner hair cells (IHC) and auditory-nerve (AN) fibers (Kujawa & Liberman, 2009), and constitutes a recently discovered form of sensorineural hearing loss besides the well-known outer and-inner hair cell damage, OHC & IHC). CS cannot be detected using the clinical audiogram or otoacoustic emissions and is therefore often referred to as “hidden” hearing loss but compromises supra-threshold temporal coding (Shaheen et al., 2015; Parthasarathy and Kujawa, 2018). Over the past ∼15 years, animal studies have provided substantial knowledge about the mechanisms of CS, showing that it develops with the normal aging process, and, strikingly, that it can also occur after temporary noise exposure without associated OHC damage (Liberman & Liberman, 2015). Computational studies simulating this deficit shows that it indeed strongly impacts the quality of the neural signal sent from the periphery to higher auditory stages (Bharadwaj et al., 2014; Verhulst et al., 2018; Lesica, 2018; Vasilikov et al., 2021). Yet, recent studies investigating the conditions and the extent to which CS affects the processing of supra-threshold sounds in humans, as involved in SPiN understanding, remain at their premises, and their outcomes are still mixed (Liberman et al., 2016; Bramhall et al., 2019; Guest et al., 2019, Prendergast et al., 2019, Garret et al., 2024). A second problem in connecting CS to sound perception deficits relates to the impossibility to quantify CS directly in humans. Through the combined use of animal models of CS and computational models of human generators of auditory evoked potentials such as the auditory brainstem response (e.g., Verhulst et al., 2016, 2018; Encina-llamas et al., 2019; Vasilkov et al., 2021; Märcher-Rørsted et al., 2022), several indirect electrophysiological markers of CS have been proposed.

Notably, the frequency following response (FFR) to sustained, sometimes modulated stimuli, reflects the fidelity of neural AN or brainstem encoding of temporal information (Bidelman et al., 2018; Krizman and Kraus, 2019). Most of these studies consider the envelope component of the FFR, referred to as FFR_env_, or the *envelope following response* (EFR). Garrett (2024) and Mepani et al. (2021) analyzed the FFR_env_ to a rectangular amplitude-modulated (RAM) 4-kHz tone. This stimulus was designed using a computational model as a specific marker of CS (Verhulst et al., 2018, Vasilkov et al., 2021). In a large cohort of 20-60 y.o. participants with clinically-normal audiograms, significant correlations between the FFR_env_ magnitude and SPiN performance were found (Mepani et al., 2021). Similar conclusions regarding the FFR_env_ in older listeners with normal audiograms were drawn by Garrett (2024), wherein both studies interpreted the relation between between FFR_env_ of the RAM tone and SPiN performance as reflecting the extent to which CS compromises envelope coding fidelity and thereby contributes to SPiN variability. A second branch of FFRs focuses on the temporal-fine structure (TFS) component of the FFR recorded to vowel sounds with formants (FFR_tfs_). TFS refers to the fast time-varying information contained within evoking sound and is neurally encoded for temporal fluctuations up to 1-2 kHz, where AN phase-locking is possible (Moore, 2007). In response to vowel-like sounds, the FFR_tfs_ exhibit a harmonic structure with highest peaks corresponding to resolved harmonics of in the input waveform, notably the first two formant peaks (Aiken & Picton, 2008; Ananthakrishnan et al., 2016, 2017). Previous works mainly studied effects of aging and hearing-loss on FFR_tfs_ were reduced FFR_tfs_ strength was observed in older compared to younger individuals with normal NH thresholds in the frequency range of the stimuli. Similar trends were seen in response to static or dynamic pure tones (Clinard et al., 2010; Clinard and Cotter, 2015; Märcher-Rørsted et al., 2022), as well as to speech-like sounds (Anderson et al., 2012, Mamo et al., 2016). FFR_tfs_ to vowel sounds were also found to be smaller in HI compared to NH participants (Ananthakrishnan et al., 2016; Molis et al. 2023). Although it is suggested age- or hearing-loss TFS-coding decline, as reflected in the FFR_tfs_, may contribute to poorer speech understanding, this relationship was not tested empirically in these studies. At the same time, further work is needed to develop a viable low-frequency FFR_tfs_ counterpart to the FFR_env_ derived with the RAM tone reflecting high-frequency processing, as it is well known that TFS based auditory cues play a pivotal role for SPiN understanding (Drullman 1995; Lorenzi et al., 2006; Parthasarathy et al., 2020).

To provide a better understanding on the triangular relationship between electrophysiological FFR markers and S(T)M-based psychoacoustic markers of supra-threshold temporal processing this study takes an integrated approach to study their interrelationship as well as the connection to CS and speech intelligibility deficits. We addressed the relevance of SM stimuli to assess TFS-coding fidelity and suggest novel psychophysical and electrophysiological markers of CS based on phase-locking capacities in the low frequencies and complemented these measurements with computational modeling. We considered two groups of young (∼20 y.o) and older (∼ 50 y.o.) individuals with clinically-normal audiograms, and a third age-matched older group with clinical hearing loss. We hypothesize that a main effect of age-related CS is observed between the younger and older normal-hearing groups, and that a main effects of OHC damage is observed between the older normal and hearing-impaired groups. We tested the extent to which our psychophysical and electrophysiological measurements were able to predict speech-in-noise performance variability measured in all individuals, specifically under a low-pass frequency filtering condition aiming to enhance the role of TFS cues. Our results suggest that FFRs to SM signals provide a novel tool to monitor CS and can be used to help teasing apart the exact origins of speech-in-noise deficits that cannot be explained using standard audiometric hearing thresholds.

## Material and Methods

### Participants and initial hearing screening

Forty-nine subjects were initially recruited in our study. These subjects were divided into three groups depending on their age and pure-tone audiogram, as described below: young normal-hearing (yNH), older normal-hearing (oNH) and older hearing-impaired (oHI). The age range of recruitment of participants was 20-25 for the yNH group, and 40-60 for oNH and oHI groups. Hearing status was first self-assessed by the subjects using a questionnaire where we used the following exclusion criteria: (i) chronic tinnitus or hyperacusis, (ii) middle-ear pathologies and/or history of middle-ear surgery, (iii) pregnancy and (iv) any known genetic hearing loss. A clinical evaluation of the participants’ hearing status was then conducted using tympanometry and otoscopy to more formally exclude cases of conductive hearing losses and/or middle/outer ear pathologies. Pure-tone audiometry was conducted at conventional frequencies (0.125, 0.25, 0.5, 1, 2, 3, 4, 6 and 8 kHz) using an Equinox Interacoustics audiometer with Interacoustics TDH-39 headphones and also at extended high frequencies (10, 12.5, 14, 16 kHz) using circumoral Sennheiser HAD-200 headphones. For participants with air-conduction thresholds exceeding 20 dB HL between 0.25 and 4 kHz, bone conduction was also measured using an Interacoustics bone vibrator placed on the mastoid to exclude potential middle-ear pathology (air-bone gaps were all < 15 dB HL).

Individuals were considered as normal-hearing if their threshold between 0.125 and 4 kHz did not exceed 20 dB HL, and thus assigned to the yNH or oNH groups depending on their age (see above). The inclusion criterion for individuals in the oHI group was a high-frequency sloping bilateral hearing loss with a threshold higher than 20 dB HL at 4000 Hz. Because several subjects clearly showed unreliable ABRs or EFRs/FFRs, we had to exclude one participant from the oHI group, two participants from the oNH group and one participant from the yNH group. In the end, the present study was made of the yNH group (n= 15, 12 females, age 21 ± 1), the oNH group (n= 16, 11 females, age 47 ± 6) and the oHI group (n= 14, 8 females, age 52 ± 6). The audiograms of the individuals of these three groups are plotted in Figure 2A.

**Figure 2.**
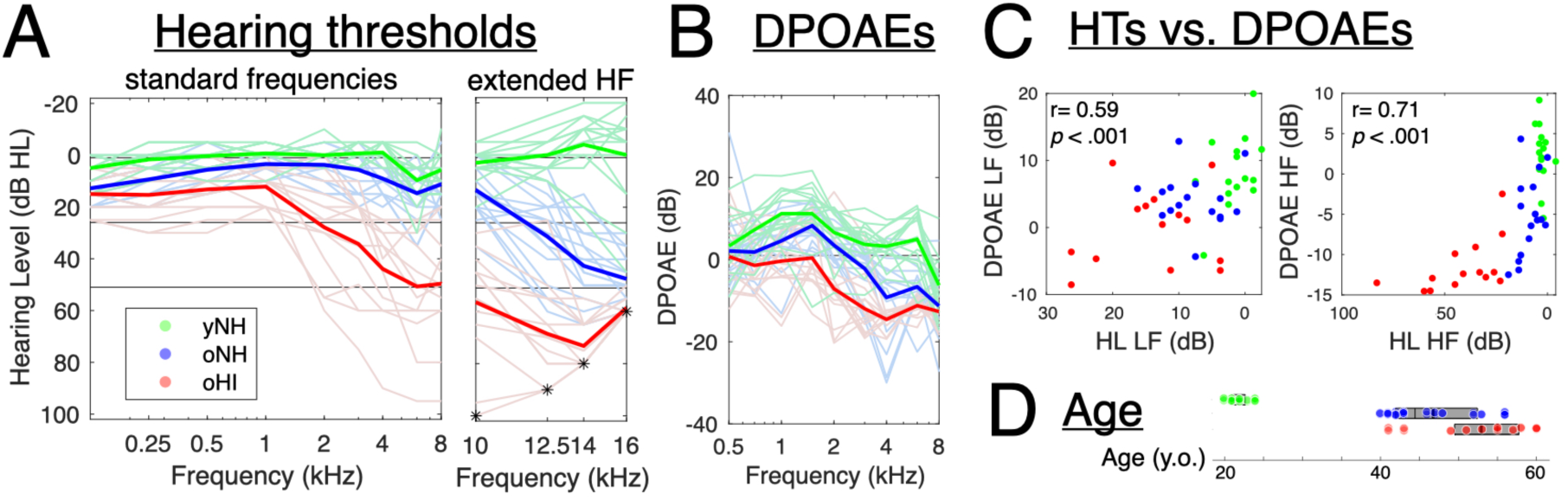
(A) Pure-tone audiograms (PTA) in standard frequencies and extended high-frequency (yNH group in green, oNH group in blue, oHI group in red); stars in the extended high-frequency range show maximum presentation levels permitted with the experimental setup. (B) DPOAEs for the three groups of participants. (C) Relationship between hearing thresholds (HTs, expressed in hearing levels, HL) and DPOAEs either in the low-frequency (LF) range (left panel) or in the high-frequency (HF) range (right panel) obtained averaging the measurements below 1kHz or above 2kHz, respectively. (D) Age of individuals in the three groups.

The study was approved by the ethical commission of Ghent University Hospital (UZ Ghent protocol number BC-08340 E01) and was conducted in accordance with the Declaration of Helsinki. All subjects were informed about the experimental procedures, provided written informed consent, and received financial compensation for their participation.

### Distortion product of otoacoustic emissions (DPOAEs)

OAEs were measured to assess OHC-integrity. We used the Universal Smart Box (Intelligent Hearing System) with Etymotic ER10D probes and analyzed with the corresponding software (SmartDPOAE and SmartEPCAM). For the DPgram, two tone pairs were presented simultaneously with f2=1.22*f1 and f2 ranging from 500 to 8000 Hz with 2 freqs/octave. The intensities were kept at 65 and 55 dB SPL for L1 and L2 respectively in a first run, (a second run was conducted with L1 and L2 at 55 and 40 dB SPL, respectively, but the results are not included in the present paper). 32 sweeps were taken and the SNR on each frequency was used for further analysis. DPOAE input/output thresholds were also measured at 1, 2 and 4 kHz as described by Verhulst et al. (2016) but are not included in the present analysis. Tone pairs were presented at f2=1.22*f1. The intensity of L2 ranged from 35 to 70 dB SPL and L1 was computed based on the Scissors paradigm of Kummer et al. (1998) (L1=0.4*L2+39). For TEOAEs, clicks of 80 µs were produced at 80 dB SPL during 2000 sweeps but were not included neither. These values were finally noise-floor corrected, i.e. by replacing them with the noise floor amplitude when lower.

### Psychophysical assessment of S(T)M sensitivity

#### Stimuli

Three types of ripple stimuli were used, with parameters derived from previous psychoacoustical studies (Bernstein et al., 2013; 2016; Miller et al., 2018; Zaar et al., 2023a). We tested different SM configurations with both noise and harmonic carriers, as well STM with harmonic carriers, and hypothesized that they would lead to comparable results (Zaar et al., 2023). We used the same technique as described in details in recent studies (Zaar et al., 2023a) to synthesize these types of stimuli, so we only report here the main aspects of this synthesis technique as well as the specific parameters that were used in the present study. Briefly, these stimuli are similarly synthesized using band-restricted carriers that are then multiplied by a modulator to impose a ripple structure on their envelope. The carrier is made of log-spaced sinusoids with random phases, with spacing on the frequency axis either extremely dense, resulting in a pink noise carrier, or following a harmonic structure to create a complex tone with a given fundamental frequency f_0_. The modulator is defined in the spectro-temporal domain, and is used to imposed either spectral modulations only or spectro-temporal modulations (STMs) on the envelope of the stimuli. The parameters of the modulator are: the modulation depth (parameter m, ranging from 0 to 1, corresponding to unmodulated to fully modulated), the phase (in radians), the scale or density (in cycles/octave, or c/o), and the rate or frequency (in Hz). In the present study, all the stimuli were 500-ms long (including 20-ms ramps at onset and offset) and had scales at 2 c/o, they varied depending on the type carrier (noise/harmonic) and the absence/presence of a temporal modulation. Importantly for the present study, the phase of the modulator was fixed at 2*pi/3, which was required because our goal was to assess the relationship between perceptual sensitivity and electrophysiological response, the latter requiring the repetition of the exact same stimulus over many trials. The parameters specific to each type of stimulus are reported below.

#### SM_noise_: spectral modulation with noise carrier

The SM_noise_ was constructed with noise carrier made by adding 400 random-phase, log-spaced sinusoids in the range [300-1500 Hz]. The modulator was only carrying spectral modulations with a scale of 2 c/o; the rate was 0 Hz.

#### SM_harm_: spectral modulation with harmonic carrier

The SM_harm_ was constructed with a harmonic carrier made by adding 13 random-phase sinusoids spaced every 110 Hz in the similar [300-1500 Hz] frequency range as for the SM_noise_ (i.e. starting at 330 Hz up to 1430 Hz), resulting in a complex tone with a virtual fundamental frequency f_0_ of 110 Hz. The modulator was only carrying spectral modulations with a scale of 2 c/o; the rate was 0 Hz.

#### STM_harm_: spectro-temporal modulation with harmonic carrier

The STM_harm_ was constructed exactly as SM_harm_, except that the modulator carrying spectral modulations with a scale of 2 c/o and also temporal modulations at a rate of 4 Hz. The 4 Hz value was chosen in light of previous studies (Bernstein et al., 2016; Miller et al. 2018) who found that the sensitivity to 2-c/o, 4-Hz STMs showed maximal correlations with SRTs. Bands of noise were also added below and above the [300-1500 Hz] frequency range of the carrier in order to mask potential temporal off-frequency cues that could be created by the temporal modulation (Narne et al., 2016). These noises were made by adding 100, random-phase, log-spaced sinusoids in the [200-300 Hz] (i.e. below) and [1500-1800 Hz] (i.e. above) regions, with levels 15 dB below the level of the sinusoids used to create the carrier.

#### Procedure

An adaptive measurement procedure similar to previous literature on this topic was used to measure ripple sensitivity for the three types of S(T)M stimuli described above. Our reasoning regarding the choice of the tracking parameters was similar to Zaar et al. (2023): our approach was to build a task that would not be too difficult and would provide fast and reliable estimates of thresholds, especially for listeners with hearing loss.

A three-interval two-alternative forced choice (3I-2AFC) procedure, where the modulation depth of the modulator was changed following a two-down, one-up procedure (Zaar et al., 2023). In all trials, the first stimulus was always unmodulated as a reference for the participant, and the modulated stimulus was presented either in the second or third interval with equal probability. Participants were instructed to select the stimulus (2^nd^ or 3^rd^) that was different from the other two. The interstimulus interval was 300 ms. Within a given trial, the starting phases of the carrier components were identical (« frozen ») across the three intervals such that the only difference was the imposed spectro-temporal modulation on the carrier; these values were refreshed from trial to trial by drawing values from a uniform distribution [0, 2π]. The modulation depth value was here defined in decibels full scale (0 dB FS) simply by taking 20 log10 (m), where m is the linear depth; 0 dB FS thus corresponds to a fully modulated stimulus. As in Miller et al. (2018), we used an initial step size of 6 dB, which was reduced to 4 dB after the first reversal for the next two reversals, and finally reduced to 2 dB until 6 reversals were reached. The thresholds used for the analysis were not those returned by an average of the modulation depth over the last reversals (see below).

Two interleaved tracks were collected per condition, and an additional third track was run if the difference between the thresholds measured (calculated here as the average modulation depth across these last 6 reversals) in the two tracks was greater than 3 dB. The modulation depth value was always limited to 0 dB FS. If the tracks did not converge because a given listener was unable to reach performance threshold by producing three consecutive incorrect responses at full modulation depth the procedure stopped and a novel experiment started using a constant stimulus procedure; this happened on 4 conditions for subjects from the oNH group (corresponding to ∼8% of measurements) and 5 conditions for subjects from the oHI group (corresponding to ∼12% of the measurements). This constant stimulus procedure was made of 60 trials presented in random order: 20 trials at full modulation depth i.e. 0 dB FS, 20 trials with modulation depth of -3 dB FS, and 20 trials with modulation depth of -6 dB FS. The threshold was then inferred from the responses collected to all these stimuli (see below).

In all cases, the thresholds were inferred from all the responses collected using a psychometric function fitting approach (detailed below) for two reasons; first because it allows us processing the data collected using a common approach for the adaptive and constant stimuli procedures, and second because we noticed after a detailed, visual inspection of the data of individual tracks with the adaptive procedure that some subjects exhibited large fluctuations around threshold, where the modulation depth first seemed to reach the threshold value, and then started to increase, which might result in poor threshold estimates. Although the reasons of why this happened were surprising to us and still remain unclear, it is an issue that could be related to task difficulty, or to the difficulty of deploying a constant attention over a certain amount time for some older and/or impaired individuals.

Because such fluctuations could lead to poorer estimates of underlying thresholds (a lack of precision that is important to address given that our goal was to examine the relationship between psychophysical thresholds and EEG data), we retained a method that, by allowing to reconstruct the whole psychometric function from all responses and then infer the threshold from this function, would be less affected by potential lapses of attention compared to a simple averaging of the modulation depth value over the final reversals. We therefore estimated the thresholds (in dB FS) corresponding to 75% correct responses after reconstruction of the whole psychometric functions for condition (using *psignifit* routine developed by Whichmann and Hill, 2001), regardless of the measurement procedure (adaptive or constant stimuli) used. A threshold was thus obtained for each individual and condition. We verified that replacing these thresholds inferred from the psychometric function by those returned by the adaptive procedure after simply averaging the last 6 reversals (an estimation of the 70.7% point on the psychometric function) yielded comparable results and would have led to similar conclusions.

All participants were tested identically on a dedicated laptop in a sound-proof room at UGent hospital (UZ). Stimuli were built and presented using an in-house Matlab program, converted through a professional soundcard (RME Fireface UCX) and delivered through headphones (Sennheiser HAD-300). Responses were entered through keyboard arrows (left arrow for 2^nd^ interval, right arrow for 3^rd^ interval). The whole experimental setup was calibrated using a Brüel & Kjær 2238 Mediator sound-level meter, coupled with the mounting plate provided for circumaural headphones. For each participant, stimuli were presented monaurally to the same ear as for the other experiments. The stimulus presented in each interval was always normalized in level and presented at 75 dB SPL (rms), i.e. there was level roving. Participants with hearing impairment were tested unaided, without their hearing aids. Measurement of sensitivity for the three types of stimuli (SM_noise_, SM_harm_, STM_harm_) was made in three separate conditions, which were conducted in a random order for each participant. A short training was provided at the beginning of the task for each of the three stimulus conditions, with longer stimuli (1000 ms) and only large modulation depths (either full modulation depth of -6dB). During the whole experiment, participants received visual trial-by-trial feedback regarding the correctness of their response, to ensure maximal engagement of attention in the task.

### Assessment of speech-in-noise intelligibility

Previous works highlighted a significant relationship between S(T)M detection thresholds and speech-reception thresholds (SRTs) in simple SPiN tasks (Bernstein et al., 2016; Miller et al., 2018). We therefore chose a standard monaural SPiN task based on a Matrix test with speech-shaped noise (SSN). Speech-in-noise intelligibility was assessed across three conditions, an unaltered, broadband (BB) condition, as well as in two frequency filtering, low-pass (LP) and high-pass (HP) conditions, detailed below. These additional conditions were designed to better tease apart the contribution of TFS and ENV cues. We reasoned that in the LP condition, participants could use both TFS and ENV cues whereas in the HP condition, only ENV cues could be used.

#### Stimuli and procedure

Speech intelligibility was measured using the Flemish Matrix sentence test (Luts et al., 2014), which consists of a closed set of non-predictable five-word sentences (10 names, 10 verbs, 10 numerals, 10 colors and 10 objects) pronounced by a single speaker. A different wordlist was used for each condition, but the same lists were used for all participants. Participants’ task was to click on the words they recognized among the list of all possible entries from the word-matrix presented on the computer screen to measure their speech reception thresholds. The speech-shaped noise had a fixed intensity of 75 dB SPL in all the conditions described below. The speech level was varied (starting at noise level -4 dB) using a one-up, one-down adaptive procedure to track the speech reception thresholds corresponding to 50% of correct word recognition. Initial step size was 5 dB and was then decreased depending on the number of correct words in each trial, reaching values smaller than 1 dB in the last reversal. Thresholds were measured in three different frequency filtering conditions; and the same filtering was used for both the speech and noise material: broadband/BB (no filtering), low-pass/LP (cutoff at 1500 Hz) and high-pass/HP (cutoff at 1650 Hz). The noise always started 500 ms before and ended 500 ms after each sentence. We initially included an additional condition to measure of speech reception thresholds in quiet (SPiQ) in a LP filtering condition, but the starting level of the speech signal (75 dB SPL) was high and did not allow measurements to converge to the threshold of 50% of correct responses within the limit of 20 trials for a few individuals having low thresholds; results from this condition are therefore not considered in the subsequent analyses. Three participants from the oHI who had the highest (worst) hearing thresholds in the high frequencies were actually not able to perceive the noise in the HP condition, and therefore their SRTs rather reflect HP speech reception thresholds in quiet and not in noise; these subjects are highlighted in Figure 3 and following analyses were thus conducted both with and without these three subjects to check for the robustness of our results. The conditions were presented in different blocks; the BB condition was always presented first, followed by the LP and HP conditions whose order was randomized across participants. For each condition, measurement was made using two interleaved tracks. A short training block was used to familiarize participants with the task.

**Figure 3.**
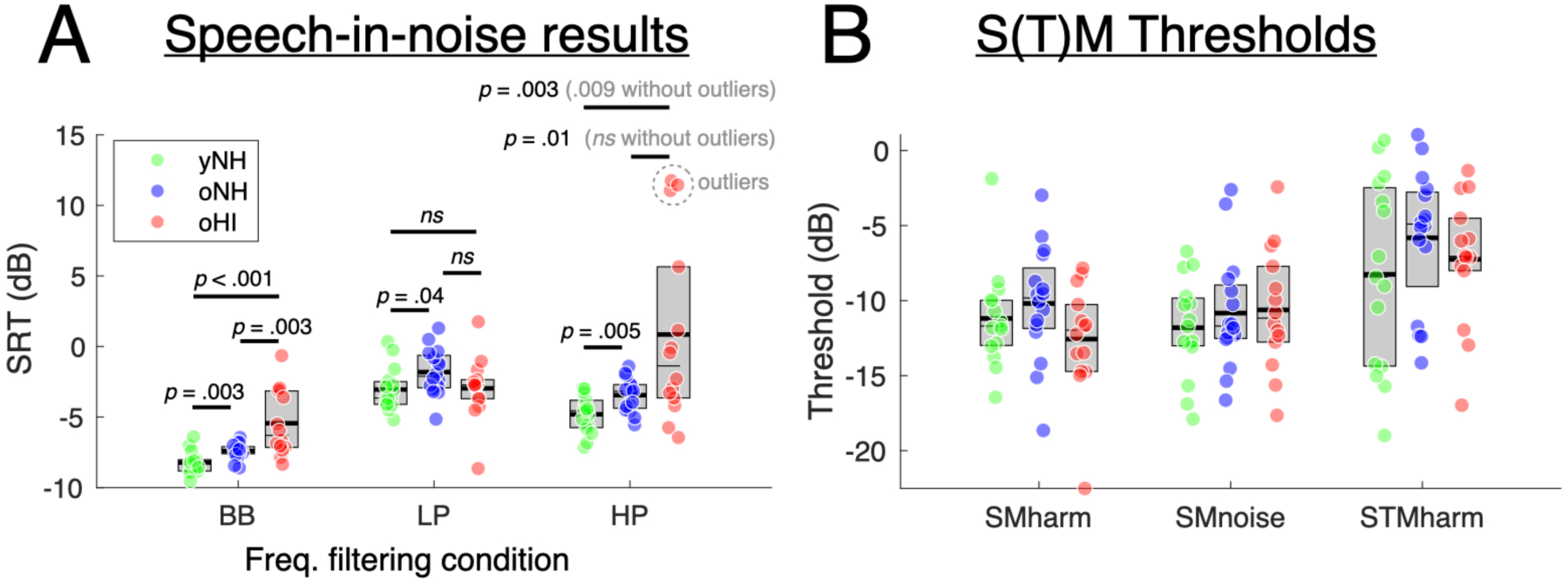
(A) Speech reception thresholds (SRTs) obtained for the three groups of listeners in the three frequency filtering conditions (broadband/intact: BB; low-pass: LP; high-pass: HP). Outliers from the oHI group in the HP conditions (see the text for details) are highlighted. P-values (uncorrected) from post hoc two-tailed t-tests conducted across groups are reported for each condition. (B) Psychophysical detection thresholds obtained for the three groups of listeners against the three types of S(T)M stimuli. No significant differences (Ps > .05) were found between groups for each type of stimulus.

For each of the three noise conditions considered for the analyses, SRTs were computed using the mean SNR/level over the last six reversals of each track, and averaged over the two tracks. The experimental setup was the same as for the measurement of STM sensitivity described above. Importantly, the stimuli were delivered monaurally to each participant in the same ear as the one used for the other tests.

### EEG measurements

#### EEG protocol and data acquisition

Measurements were conducted using the Universal Smart Box (Intelligent Hearing System) with the SmartEP continuous acquisition module (SEPCAM). Participants were seated in a reclining armchair in a quiet room, watching a silent movie in order to keep the participant relaxed and awake (Purcell et al., 2004). Ambu Neuroline snap electrodes were connected to vertex, nose and both mastoids after applying NuPrep® gel with a cotton swab for reducing the skin-impedance to a maximum of 3 kΩ. The sampling frequency was 10 kHz. The stimuli were presented with ER-2 inserts (Etymotic Research) and the ears were covered with earmuffs to minimize the noise intrusion level. All electrical devices other than the measurement equipment were turned off and unplugged.

#### Auditory brainstem response (ABR)

ABR-responses were measured to obtain an extensive representation of the auditory nerve function. The used parameters are based on the studies of Liberman et al. (2016) and Vasilkov et al. (2021). A click-stimulus of 100 µs was presented with a rate of 11 Hz. Two intensities (75 and 90 dB peSPL (rms)) were measured, with both 3000 alternating sweeps, filtered between 10 and 1500 Hz. Positive and negative peaks were pointed on the ABR-curves manually by a professional audiologist. Both amplitudes and latencies of selected peaks were extracted from these data. ABR results are not detailed in the present paper; they were primarily included to verify the reliability of our EEG measurements based on previous data from our group collected using a similar protocol. In total, four participants were excluded from the analysis because they showed unreliable ABRs (see Participants section).

#### Frequency following response (FFR) with the RAM stimulus

The RAM stimulus was used to specifically assess ENV-coding fidelity (Vasilkov et al., 2021). It uses a high-frequency pure tone carrier at 4 kHz that was 100% rectangularly amplitude modulated at 110 Hz and with a duty cycle of 25% (RAM25) as described in Vasilkov et al. (2021). The stimuli were presented at 75 dB SPL (rms) for 200 ms during 1000 alternating sweeps, and the signal was filtered between 10 and 5000 Hz.

#### Frequency following response (FFR) with the SM_harm_ stimulus

The SM_harm_ stimuli were used to specifically assess TFS-coding fidelity. Two SM_harm_ stimuli identical to those used for the psychophysical measurement; one with a 100% modulation depth (0 dB FS) and another one with a 50% modulation depth (i.e. – 6 dB FS). These stimuli were 400-ms long, had a level 75 dB SPL (rms), and were each presented using 3000 repetitions of alternating polarities monaurally to the participant’s best ear. The initial idea of using a stimulus with a 50% modulation, in addition to the stimulus with full modulation depth, was to see whether it could provide a contrast measure and therefore allow for a better assessment of the relationship with psychophysical modulation detection thresholds. Yet, we found no measurable and consistent differences between the responses to these sounds, so the responses collected with these two stimuli were aggregated to derive an average FFR for analysis.

#### FFR pre-processing and analysis

All recordings were first band-pass filtered between 60 and 2000 Hz, in order to highlight the frequency range of interest for subcortical responses induced by our stimuli. The same pipeline was used to analyze the responses to the RAM and SM_harm_ stimuli. For each stimulus type, we considered the responses recorded over the total duration for analysis. We first performed baseline correction by removing the mean computed over all responses. We then performed epoch rejection using a sliding window; the range of the signal (max – min) was extracted from each epoch, and an epoch was rejected when its range exceeded 3 times the median value computed over 200 epochs. For the RAM stimuli, an average number of 945 ± 36 (M±SD) trials were available for each participant for further analysis. For the SM_harm_ stimuli (after aggregating the responses to the two modulation depths), an average number of 2848 ± 82 trials were available for each participant for further analysis.

We then followed the now widely adopted methodology for extracting the TFS and the ENV components from FFRs (Aiken & Picton, 2008; Ananthakrishnan et al., 2016), where FFR_env_ (sometimes also referred to as envelope following response, or EFR, in other studies) and FFR_tfs_ are computed using the sum or difference of responses to positive and negative stimulus polarities, respectively. Although some works showed that these FFRs are corrupted by cochlear microphonics (Lucchetti et al., 2018; Borijin et al.,2022), if present, these effects would be similar across groups and would thus not explain the effects reported in that study. We then used a bootstrap-based correction algorithm to estimate the frequency spectra after noise-floor correction (Zhu et al., 2013, but see Vasilikov et al., 2021). Finally, a scalar value representing the overall magnitude of the responses (later used for statistical analyses) was computed by extracting and summing the magnitude of the harmonics (multiples of f0=110 Hz) that were significantly above noise-floor; we retained harmonics #1 to #7 for both the RAM stimulus and the SM_harm_ stimulus.

### Computational simulations with a model of the auditory periphery

We used a biophysically-inspired model of the auditory periphery (Verhulst et al., 2018) to simulate auditory nerve (AN) responses across the different types of fibers with high, medium and low spontaneous rates (H, M, L) to the exact same SM_harm_ stimulus as for the FFR empirical characterization, i.e. with an input level of 75 dB SPL. AN responses were extracted to both positive and negative stimulus polarities, and either summed or subtracted (across all different types of fibers) to assess the ENV and TFS part of the Wave 1 response, respectively, exactly as for the FFR analyses. A FFT of the resulting response was then computed to derive its frequency spectrum, and the first seven harmonics were averaged. Different hearing profiles were tested to simulate the hearing characteristics of the three groups of participants engaged in the present study; we tested 3×3 combinations of OHC loss (no OHC loss; normal audiometric thresholds ∼ up to 1-kHz, and then sloping audiogram above; flat audiometric thresholds ∼30 dB HL up to 1-kHz, and then sloping audiogram above) and synaptopathy (number of H/M/LSR fibers at each CF: (13/3/3), (7/0/0), (4,0,0)).

### Statistical analyses

All statistical analyses were performed using R Statistical Software (v4.3.1; R Core Team 2022). Mixed analyses of variance (mixedANOVA) were conducted using a univariate approach with Huynh-Feldt correction for the degrees of freedom (Huynh & Feldt, 1976); the correction factor 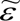 is reported, and partial 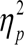 is reported as measure of association strength. Two-tailed t-tests were conducted as post hoc tests. Correlations were analyzed using Pearson correlation coefficient (r) and are reported in the text and the figures as uncorrected. Most correlations, unless mentioned otherwise, were computed after aggregating all individuals for the three groups (N=45). The significance level for all tests was set at *α* = 0.05; correction for multiple comparisons (Bonferroni) was considered for discussing the robustness of the effects when required. Multiple linear regressions were conducted using the lm function in R.

## Results

### Hearing thresholds and OAEs

Figure 2 shows hearing sensitivity assessed at threshold using pure tone audiometry (PTA, Fig. 2A), DPOAEs (Fig. 2B) and the age of the individuals in our three groups (young normal-hearing, yNH; older normal-hearing, oNH; older hearing-impaired, oHI; Fig. 2C). ONH had higher hearing thresholds compared to yNH at both standard frequencies (t(29)=5.01, *p* < .001) as well as extended high frequencies (t(29)=9.31, *p* < .001). OHI had higher hearing thresholds compared to oNH at both standard frequencies (t(28)=7.66, *p* < .001) as well as extended high frequencies (t(28)=7.65, *p* < .001). The average differences in thresholds across frequencies below 1kHz, which corresponds to the frequency spectrum of the SM_harm_ stimulus, were significant between yNH and oNH (t(29)=4.51, *p* < .001) as well as between oNH and oHI (t(28)=2.70, *p* = .012). Yet, these differences remained in a relatively narrow range of 5-15 dB. As expected with our inclusion criterion, oNH and oHI groups exhibited strongest threshold differences above 1 kHz. The DPgram analysis (Fig. 2B) shows similar results. When averaged across all frequencies, DPOAE amplitudes were higher for yNH compared to oNH (t(29)=4.58, *p* < .001), and also higher for oNH compared to oHI (t(28)=4.35, *p* < .001). When restricting these comparisons to the low-frequency range (below 1kHz), these differences were just significant after correcting for multiple comparisons (n=2); (yNH vs. oNH: t(29)=2.36, *p* = .05; oNH vs. oHI: t(28)=2.43, *p* = .04). As shown in Fig. 2C, DPOAEs correlated with hearing thresholds when considering either the low-frequency (r=0.59, *p* < .001, aggregating across all N=45 individuals) or the high-frequency (r=0.71, *p* < .001, aggregating across all N=45 individuals) ranges. The DPOAEs of some individuals in the oHI group were around noise-floor level in the high-frequency range, which resulted in a flooring effect (see left panel). In sum, PTA and DPOAE measurements were consistent and strongly paralleled each other, supporting the view that both could be taken as alternative proxies of OHC damage. Together, these measurements suggest that the differences in OHC damage in the low frequencies (below 1kHz) between our three groups, if any, could be considered as rather small, and that the main differences between groups were in the high frequencies and in the extended high frequencies.

### SRTs across frequency filtering conditions

Figure 3A shows the results of the speech-in-noise tests, expressed in speech reception thresholds (SRT), for the three groups and the three frequency filtering conditions (BB, LP, HP). Descriptively, as expected, SRTs were lower (better) in the BB condition than in the LP and HP conditions where less cues are available. We conducted a mixedANOVA on the SRTs, with the group (yNH, oNH, oHI) as between-subject factor and condition (BB, LP, HP) as within-subject factor. We found a significant effect of group (*F*(2,42) = 7.58, *p* < .001, 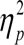 = 0.265), a significant effect of condition (*F*(2,84) = 78.92, 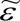= 0.632, *p* < .001, 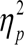 = 0.653), and a significant group x condition interaction (*F*(4,84) = 9.63, 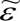 = 0.632, *p* < .001, 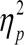 = 0.314). Post-hoc two tailed t-tests were then conducted to compare the three groups for each stimulus type; p-values (uncorrected) are reported Fig. 3A. SRTs were lower for yNH compared to oNH, and for oNH compared to oHI in BB and HP conditions, and these differences were significant (Ps < .05) after Bonferroni correction for multiple comparisons (n=3). In the HP condition, the difference between oNH and oHI was not significant after excluding the three outliers (see above), but other group differences remained significant. In comparison, there was no significant group differences in the LP condition after correction for multiple comparisons (n=3). To further address the potential factors limiting speech-in-noise understanding, we computed the correlations between the individual thresholds across these three conditions (see Fig. S1B, left panels). For the yNH group, SRTs in the BB were significantly correlated to the SRTs in both the LP (p < .001) and HP conditions (p = .002). Interestingly, these analyses revealed that for the oNH group, SRTs in the BB were significantly correlated to the SRTs in the LP condition only (p = 0.03), while in contrast, for the oHI group, SRTs in the BB were significantly correlated to the SRTs in the HP condition only (p < 0.001).

### Psychophysical detection thresholds for S(T)M sounds

Figure 3B shows the results of the psychophysical tests assessing detection thresholds for the three groups and the three types of stimuli (SM_harm_, SM_noise_, STM_harm_). First, at a descriptive level, we observed a large variability across individuals in all conditions, with differences amounting 5 to 15 dB between extreme listeners, with no obvious differences between groups. Thresholds were comparable between SM_harm_ and SM_noise_, in the range of -10 to -12 dB; in comparison, they were higher (worst) for the STM_harm_ stimulus. A mixedANOVA conducted on these thresholds, with the group (yNH, oNH, oHI) as between-subject factor and stimulus type (SM_harm_, SM_noise_, STM_harm_) as within-subject factor, supported these observations. There was a significant effect of stimulus type only (*F*(2,84) = 21.39, 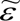 = 0.806, *p* < .001, 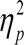 = 0.338); the effect of group was not significant (*F*(2,42) = 0.95, p=.396, 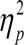 = 0.043) and the group x stimulus type interaction was also not significant (*F*(4,84) = 0.92 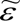 = 0.806, *p* = .456, 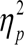 = 0.042).

To address the extent to which the same auditory processes are engaged with the different types of stimuli, we tested whether the inter-individual variability of thresholds for one stimulus type was related to the inter-individual for another stimulus type. We computed the correlations across conditions, after aggregating all individuals from the three groups together (see Fig. S2). These correlations were significant between SM_harm_ and SM_noise_ (p < .001) as well as between STM_harm_ and SM_noise_ (p = .007), but were not significant between SM_harm_ and STM_harm_(Ps > .05) These results thus only partly support our hypothesis that the thresholds measured for these different types of stimuli reflect the exact same combination of processes.

### FFRs to the RAM tone and the SM_harm_ stimulus

Figures 4 and 5 show the average ENV and TFS components of FFRs, i.e. FFR_env_ and FFR_tfs_, measured for the three groups of listeners (yNH, oNH, oHI) with the RAM tone and the SM_harm_ stimulus, respectively. For the RAM tone (Fig. 4), clear peaks are observed at various harmonics of the fundamental frequency (f_0_ = 110 Hz) in the FFR_env_ (Fig. 4A, top panels). In sharp contrast, FFR_tfs_ of the RAM tone does not carry any clear structure; peaks are in most cases not even above noise floor (Fig. 4A, bottom panels). This difference in response between FFR_env_ and FFR_tfs_ is expected because the RAM tone uses a high-frequency carrier at 4 kHz, which is far above the limit of TFS coding, so only its ENV is encoded and this is reflected in the FFR. For the SM_harm_ stimulus, peaks at multiple of f_0_ can be observed in both the FFR_env_ and FFR_tfs_; yet, it is important to note that the y-scale is reduced by a factor of 10 between Figure 4 and 5. These data for the SM_harm_ corroborate previous observations with complex tones or vowels showing that both the ENV and the TFS are neurally encoded, yielding FFR_env_ and FFR_tfs_ spectra with peaks of comparable magnitude. Also, as expected, the peaks in FFR_env_ were present for lower harmonics ((#1 to #4) for each individual and type of stimuli while the peaks in FFR_tfs_ were mostly visible at higher harmonics (#4 to #6).

**Figure 4.**
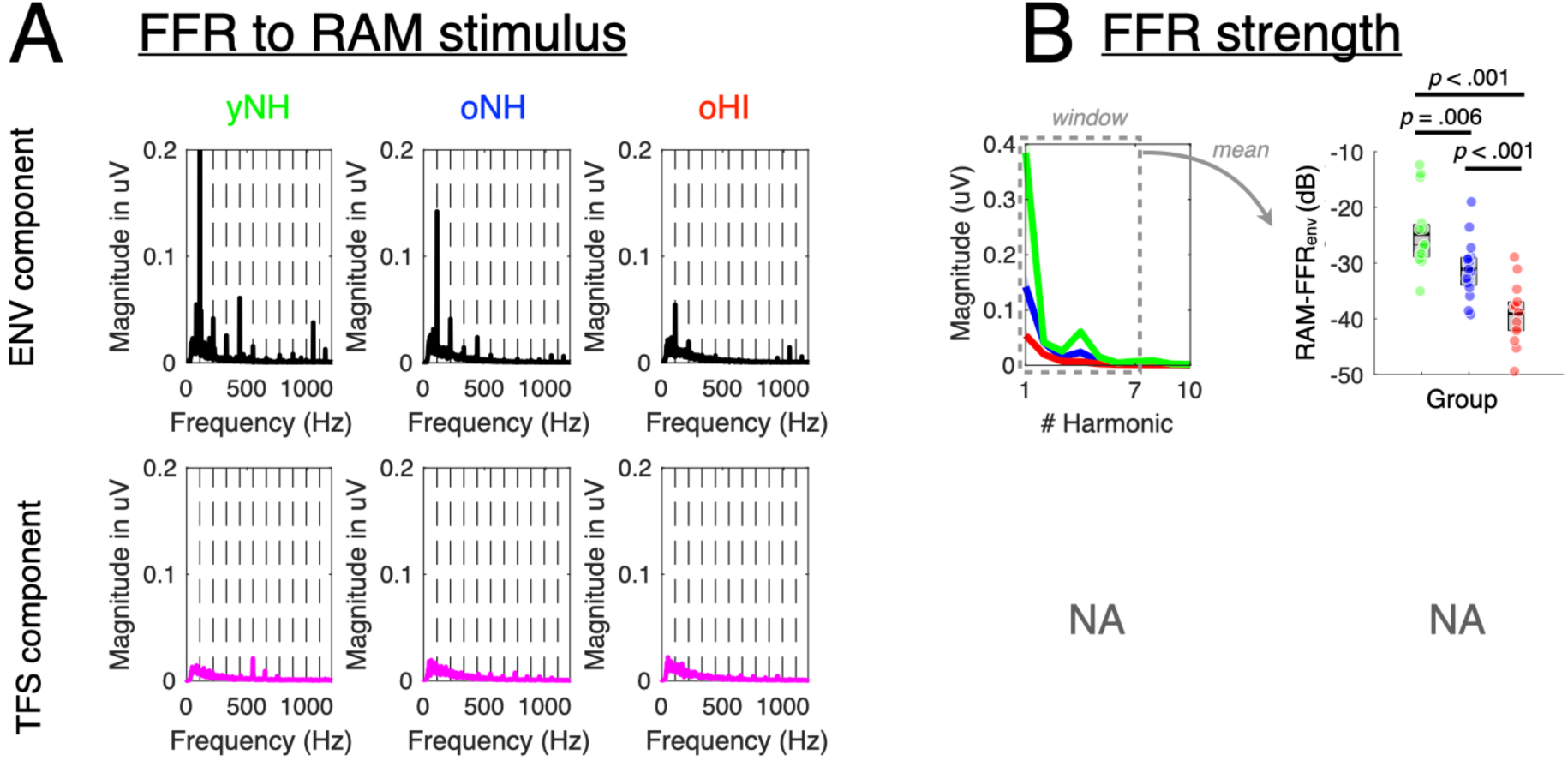
(A) Average ENV and TFS components of FFRs to the RAM stimulus obtained in the three groups of subjects (yNH, oNH and oHI) after noise-floor correction (see Methods). Peaks at multiples of f0 (110 Hz), which correspond to neural phase-locking to stimulus characteristics, align with dashed lines; peaks falling outside these lines correspond to electrical noise in the measurement system. There was a clear harmonic structure in the FFR_env_ (top), but not in the FFR_tfs_ (bottom). Note that the amplitude of the first peak (110Hz) in FFR_env_ for the yNH group extends beyond the displayed range (value is ∼ 0.4 µV, see panel B). (B) Right panels show the magnitude of harmonics #1 to #10 extracted from FFR_env_ (top) for the three groups (shaded regions, when visible, show standard error of the mean). Right panels show FFR strength for ENV component assessed by averaging the amplitude of harmonics #1 to #7, and then taking the log of this value. P-values (uncorrected) from two-tailed t-tested conducted on these scalar values expressed in dB are reported in the panel. These analyses were not conducted on FFR_tfs_ (bottom), and are referred to as NA (not applicable).

**Figure 5.**
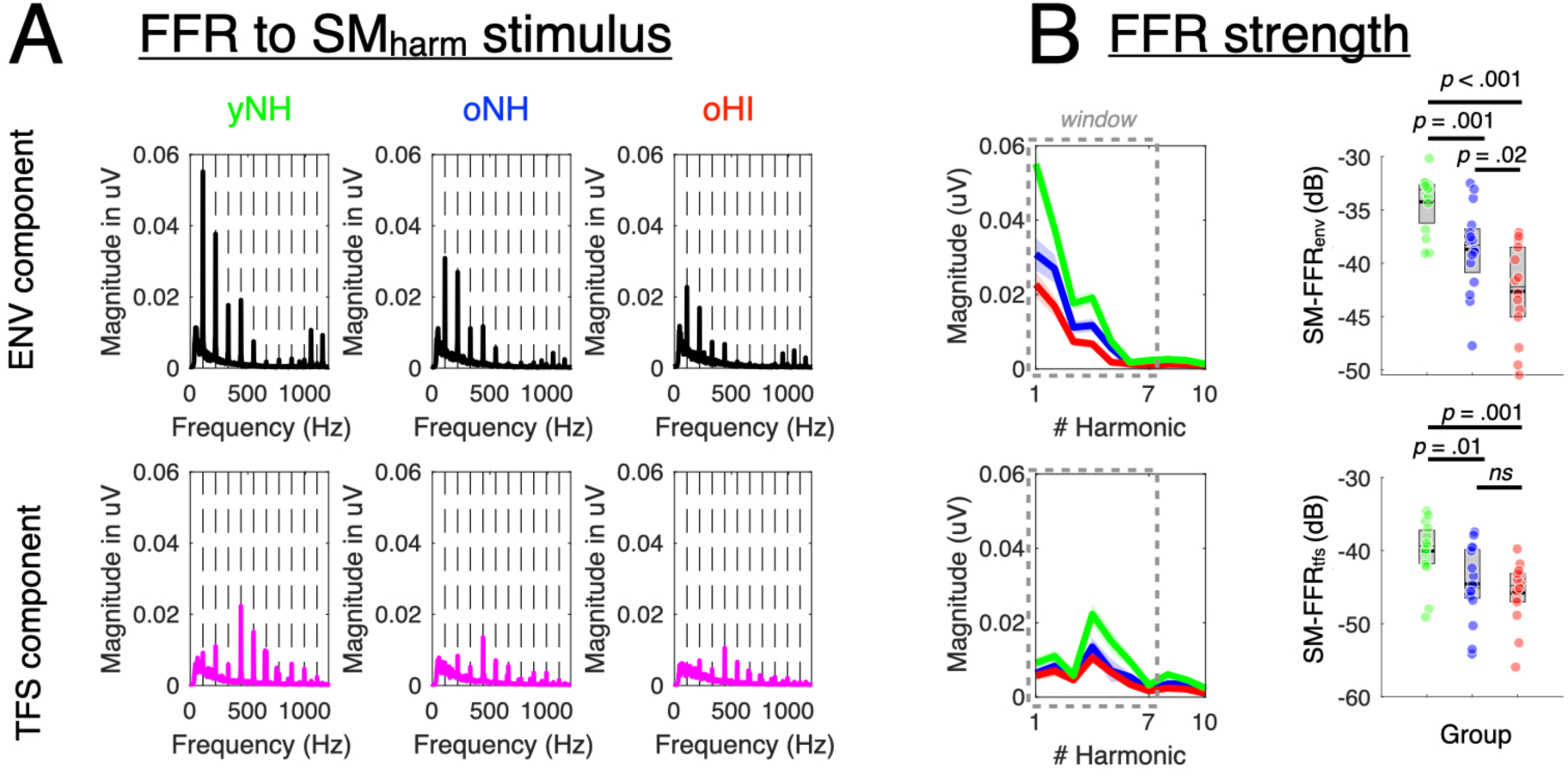
(A, B) Same as Fig. 4, for both the FFR_env_ and FFR_tfs_ to the SM_harm_ stimulus (Fig. 2B).

Descriptively, these plots show higher responses in all cases for yNH compared to oNH, and also for oNH compared to oHI listeners. To analyze these group differences more quantitatively, we followed previous approaches: we extracted the average magnitude of the seven first harmonics (which contain most of the energy related to the neural phase-locking to the stimulus ENV and TFS components) in the spectra of FFR_env_ and FFR_tfs_ to compute one single scalar value for each individual, type of stimulus, and components (ENV / TFS), except for the TFS component of the FFR obtained with the RAM tone that showed no measurable structure. This scalar, then expressed in dB, reflects the strength of coding of the ENV / TFS component and is here referred to as FFR_env_ and FFR_tfs_ strength, respectively. We considered the same harmonics #1 to #7 for analyzing the results obtained with the RAM tone and the SM_harm_, but we verified that our results were robust to other choices of harmonic numbers. The results of these analyses are presented in Figures 4B and 5B; t-tests were used to compare the three groups, and uncorrected p-values are directly reported in the panel of these figures. There was a clear primary effect of age: yNH had significantly greater FFR strength than oNH or oHI, and this effect was found considering either FFR_env_ or FFR_tfs_, and considering for both the RAM tone and the SM_harm_ stimulus. Second, there was an effect of hearing thresholds: oNH had significantly greater FFR strength than oHI, and this effect was found considering either FFR_env_ or FFR_tfs_ components obtained with the RAM tone, but not for any of the components derived with the SM_harm_ stimulus (the difference observed between oNH and oHI for the FFR component at p = .02 did not survive correction for multiple comparison, n=3).

Using correlational analyses, we examined the relationships between the scalar indexes reflecting FFR_env_ and FFR_tfs_ strength for the two types of stimuli, the RAM tone and the SM_harm_ stimulus, which primarily engage higher and lower portions of the CFs, respectively. We found that these measurements were significantly related (Fig S3). We found strong and significant correlations between the FFR_env_ obtained with the two stimuli (r=0.69, *p* < .001; Fig. S3A), and between the FFR_env_ and the FFR_tfs_ obtained with the SM_harm_ (r=0.70, *p* < .001; Fig. S3B). The correlation between the FFR_env_ obtained with the RAM tone and the FFR_tfs_ obtained with the SM_harm_ was also significant (r=0.37, *p* = .01; Fig. S3C), and appeared to be primarily driven by higher harmonics (harmonic #4 to harmonic #7; see Fig. 3C). In what follows, we only consider RAM-FFR_env_ as a marker of neural ENV coding to assess the relationship with other variables, as it corresponds to the ENV-based marker with the highest signal-to-noise ratio.

### Relationships between SRTs and psychophysical and neural markers of TFS coding

We here further addressed whether the psychophysical or electrophysiological measurements presented above would allow accounting for the variance in SPiN thresholds across listeners. In a first step, we computed simple regressions between SRTs and individual variables from our empirical measurements. We first examined the relationships between SRTs in the BB, HP and LP conditions, and FFRs (Figure 6A). For the SM_harm_ stimulus, used to assess TFS coding fidelity in the low frequencies, all correlations were not significant (Ps > .05). For the RAM tone stimulus, used to assess ENV coding fidelity in the high frequencies, the correlations between FFR_env_ and SRTs in the BB and HP conditions were significantly negative (r=-0.47, *p* = .001 and r=-0.46, *p* = .001, respectively). The interpretation of these correlations is further discussed in the next section, where other predictors such as PTA or age are included in the models. Next, we examined the relationships between SRTs in BB and LP conditions and psychophysical thresholds obtained in particular with SM_harm_ and SM_noise_ (Figure 6B) for which our measurements seem most reliable (see above). These correlations were non-significant (for SRT in the LP condition and the thresholds for the SM_noise_, the correlation was r=0.34, *p* = .021 before PTA correction, and r = 0.32; *p* = 0.03 after PTA correction, but this was not significant after correction for multiple comparisons). Finally, we analyzed the relationship between FFR_tfs_ and detection thresholds collected with the SM_harm_, both hypothesized as potential proxies of TFS coding (see Fig. 6C). We found no significant correlation between both measurements (*p* > .05)

**Figure 6.**
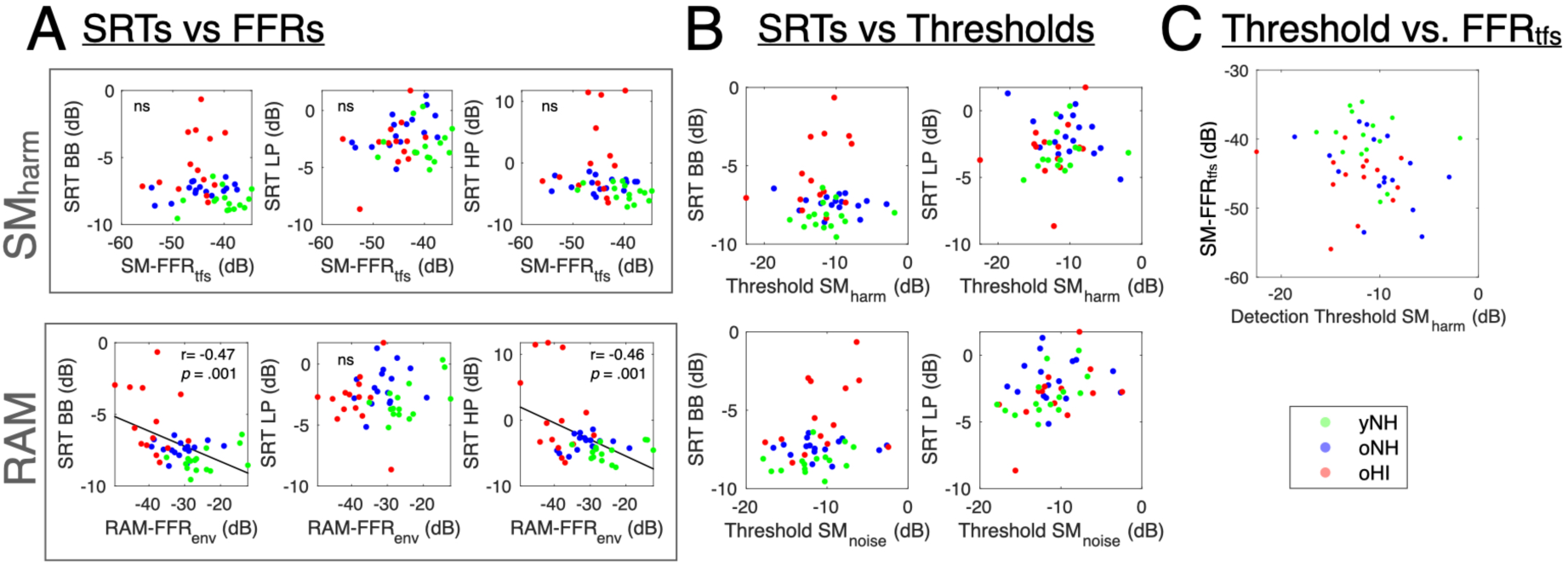
(A). Relationships between SRTs the three conditions (BB, LP, HP) and FFRs for the SM stimulus (SM-FFR_tfs_, top panels) and RAM tone (RAM-FFR_env_, bottom panels). (B) Relationships between SRTs and psychophysical detection thresholds measured with the SM_harm_ (top panels) and the SM_harm_ stimulus (bottom panels). (C) Relationship between and psychophysical detection thresholds and FFR_tfs_ (SM-FFR_tfs_) both obtained with the same SM_harm_ stimulus.

In a second step, we tested the extent to which multiple linear regression models including several predictors in addition to the FFR-based predictors could account for the interindividual variability in SRTs. We adopted a hypothesis-driven approach, starting from the full model (without interactions) that included all possible predictors, and progressively removed them one by one until we converged toward a stable model with a minimum of significant predictor(s) remaining. The outputs (estimates, p-values) of the different regressions are shown in Table S1; we only briefly summarize the main conclusions below.

The full model included five different predictors, FFR_tfs_ from the SM_harm_ stimulus (or SM-FFR FFR_tfs_), FFR_env_ from the RAM tone (or RAM-FFR_env_), hearing thresholds in the low frequencies 0.125-1 kHz (HT_LF_), hearing thresholds in the high frequencies 2-8 kHz (HT_LF_), SM_harm_ psychophysical thresholds, and age.

First, we aimed to predict SRT in the BB condition. The full model returned only HT_HF_ as a significant predictor (Est=0.08; p < .001). Yet, it is known that HT_HF_ obscures the potential effect of our predictors of synaptopathy; HT_H_ is found to enhance synaptopathy and thus be correlated with these predictors. When HT_HF_ is removed, RAM-FFR_env_ indeed became significantly negatively correlated (Est=-0.09; p =0.03). Table 1 shows that this significance hold for all nested models, when other predictors were removed one by one. The same conclusion was reached when yNH subjects were excluded. In sum, there is a robust negative correlation between SRT in the BB condition and RAM-FFR_env_. Next, we followed the exact same approach to account for SRT in the HP condition. We reached the same conclusion as with SRT in the BB condition: there was a robust negative correlation between SRT in the HP condition and RAM-FFR_env_. This correlation still held after removing other predictors one by one, and also when the three outlier subjects in this task (see above) were excluded. Thirdly and lastly, we aimed predict SRT in the LP condition. The full model returned SM-FFR_tfs_ and age as significant predictors (p < .05). Following the same reasoning, we removed age as predictor in the model, since age is known to enhance synaptopathy, and as such may obscure the true effect of other predictors. When this was done, no significant predictors remained. Age was therefore put back to the model, and the other predictors were removed one by one. Once this was done, SM-FFR_tfs_ and age became again significant predictors (p < .05), showing positive correlations with SRT. We asked whether these observations related to the age factor would hold after excluding yNH individuals from the regression, since the main age difference between our subjects is driven by individuals from that group. When doing so, age was not a significant predictor anymore, and FFR_tfs_ was the only significant predictor in all models tested.

### Relationships between age on FFR_tfs_ and FFR_env_ assessed using correlational analyses

To complement the age-related effects highlighted by the group-level comparisons (see above), we used a correlational approach to further assess the effect of age on RAM-FFRs and SM-FFR_tfs_. For these analyses, we considered the individuals from the oNH and oHI groups only to avoid any ‘bias’ induced by the lower average age of the yNH group. We did not find any significant correlation between RAM-FFR_env_ and age (*p* > 0.05; see Fig. S4). However, we found a significant correlation between SM-FFR_tfs_ and age (Fig. 7A, r=-0.47; *p* = 0.008). Because PTA and age are two intrinsically linked and highly correlated factors, we tested whether this correlation would hold after accounting for variation in PTA. To perform this correction, we first conducted a linear regression between SM-FFR_tfs_ and PTA and then assessed the relationship between the residuals from this regression, which reflect the variance in SM-FFR_tfs_ that cannot be explained by the variance in PTA with age; we found that the correlation remained significant after correcting for PTA (Fig. 7B, r=-0.45; *p* = 0.013). We then asked which specific harmonic components of the SM-FFR_tfs_ would primarily underlie this correlation. To do so, we conducted correlations between the magnitude of individual harmonics (from #1 to #7) and age, and found that only magnitude for harmonic #4 was significant (r=-0.57; *p* = 0.001), even after adjusting for PTA (r=-0.55; *p* = 0.002). This result indicated that the SM-FFR_tfs_ correlation with age was primarily driven by harmonic #4 (the only correlation that remained significant after Bonferroni correction for multiple comparisons, n=7), which incidentally also corresponds to the harmonic of highest energy contribution to the total FFR.

**Figure 7.**
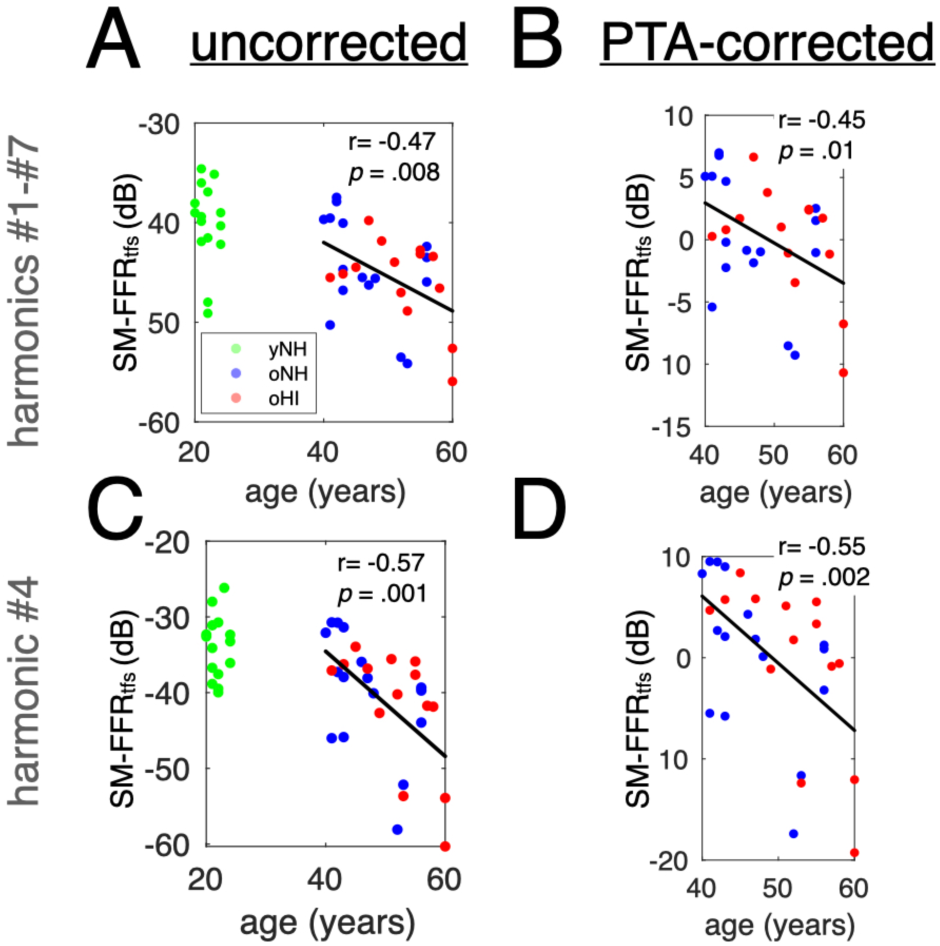
(A-D) Relationships between SM-FFR_tfs_ strength and age without correction (left panels) or after PTA-correction (right panels; see the text for details), considering either the average magnitude of harmonics #1-#7 (top panels) or harmonic #4 (bottom panels). In all panels, black lines show the linear regressions lines from analyses performed on the aggregated data of oNH and oHI individuals.

The fact that a negative correlation was observed only with the TFS-based FFR marker but not with the ENV-based FFR marker can likely be understood by considering the differences in frequencies covered by their respective carrier stimuli. Indeed, while the SM_harm_ covers lower frequencies (<1.5 kHz) at which older individuals all had comparable hearing thresholds (Fig. 2A), the RAM tone covers high frequencies where differences in hearing thresholds also contribute to the variability among oNH and oHI listeners and would presumably make the relationship with age less clear.

### Impacts of AN synaptopathy and OHC damage on FFR strength estimated from simulations with a computational model of the periphery

We run computational simulations using a biophysically-inspired modeled of the auditory periphery (Verhulst et al., 2018) to assess quantitatively the effects of OHC damage and cochlear synaptopathy on FFR derived with the SM_harm_ (modulation depth of 100%). Results are shown in Figure 8. While amounts of OHC damage consistent with the potential loss in our cohorts of listeners would lead to small increases of FFR_tfs_ strength of 1 to 3 dB, a simulation of different synaptopathy, here modeled as a partial loss of HSR fibers and total loss of MSR and LSR fibers (see the legend of Fig. 8 for details), leads to a stronger decrease of FFR_tfs_ strength in the range 7-15 dB, consistent with the effects highlighted in Figure 5B. It is important to note that very similar effects are predicted by the model on FFR_env_.

**Figure 8.**
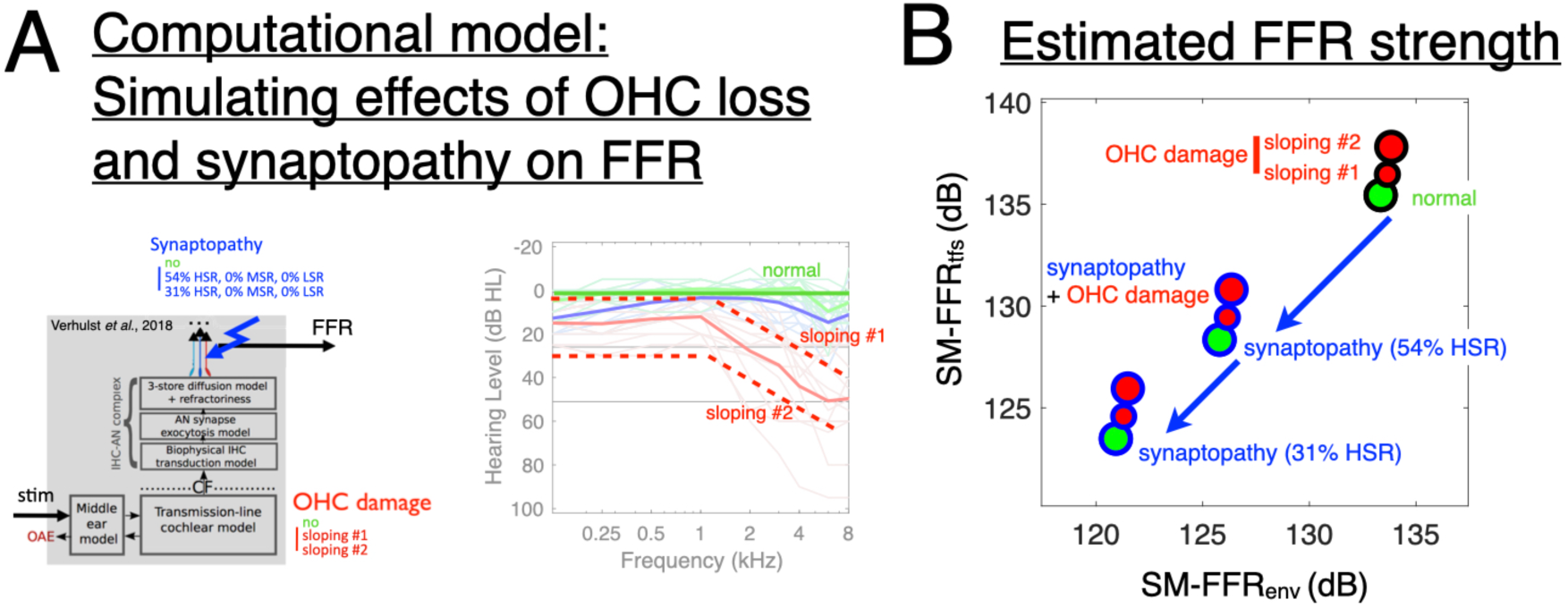
(A) Illustration of the computational model used in the present study (Verhulst *et al*., 2018) and the specific parameters adjusted for simulating different types of impairment. Model parameters were adjusted to simulate the combined effects of two aspects of hearing loss: (i) by adjusting the transmission line model coefficients to simulate different levels of OHC loss that qualitatively captured the variability of audiograms measured across our cohort of oHI listeners, as shown on the right panel where the modeled audiograms are superimposed with the measured audiograms (green: no OHC loss, red: sloping #1 and sloping #2); (ii) by reducing the number of AN fibers in the model to simulate different levels of synaptopathy (green: no synaptopathy corresponding to [13 HSR, 3 MSR, 3 LSR] AN fibers in the model, blue: [7 HSR, 0 MSR, 0 LSR] and [4 HSR, 0 MSR, 0 LSR]). The sum of AN responses across the remaining fibers was taken as an estimate of FFR. FFR_env_ and FFR_tfs_ were estimated as for the empirical work, by summing or subtracting the estimated FFRs to the stimuli with positive and negative polarities, respectively. (B) Combined effects of simulated OHC loss or/and synaptopathy estimated on FFR strength in response to the SM_harm_ stimulus used in the experiments; ENV and TFS components on the y and x axes, respectively. Marker edge colors reflect the simulation of synaptopathy in the model (black: no; blue: 54% or 31% of remaining HSR fibers) and marker face colors reflect the simulation of OHC damage (green: no; red: sloping #1 or sloping #2 audiograms). Large changes in FFR strength (7-15 dB) both on ENV and TFS components can be observed for the two levels of synaptopathy simulated, while in contrast the different levels of simulated OHC loss yield much smaller effects (1-3 dB).

## Discussion

In this study, we investigated the impact of age and hearing loss on the neural and perceptual coding of the temporal envelope (ENV) and temporal fine structure (TFS) in complex sounds. We conducted frequency-following responses (FFRs) and psychophysical measurements in three groups of listeners: young normal-hearing (yNH), older normal-hearing (oNH), and older hearing-impaired (oHI). Our primary goals were to (1) evaluate whether these measures could serve as potential proxies for cochlear synaptopathy (CS) and (2) assess their ability to predict interindividual variability in speech-in-noise (SPiN) performance. To assess ENV coding, we used a 110-Hz rectangular amplitude-modulated (RAM) tone with a 4-kHz high-frequency carrier, based on the methodology of Vasilikov et al. (2021). For TFS coding, we employed a low-frequency (300-1500 Hz), harmonic complex tone (f_0_= 110 Hz) with spectral modulations (SM_harm_), a stimulus derived from psychoacoustic studies on supra-threshold hearing deficits (Bernstein et al., 2016; Zaar et al., 2023a). This dual approach allowed us to assess both ENV and TFS coding at the electrophysiological (FFR) level and via psychophysical modulation detection thresholds. Speech-in-noise performance was evaluated in three conditions: unaltered broadband (BB) stimuli, low-pass (LP) filtered stimuli, and high-pass (HP) filtered stimuli. The LP and HP conditions were included to compare the relative contributions of ENV vs. TFS cues in speech perception.

### SRTs across low-pass and high-pass frequency filtering conditions point towards different sources of neural coding deficits

We used the approach of recent studies that consists in low-pass / high-pass filtering the speech and the noise material to preferentially target the frequency ranges in which the ENV / TFS coding mechanisms operate, respectively (Verhulst and Warzybock, 2018, Garrett et al., 2024). Overall, results of speech-in-noise tests (see Fig. 3) exhibited lower (better) SRTs in the BB compared to the LP and HP conditions for all groups, a trend that is expected because less cues are available in the frequency-filtered conditions compared to the intact (unfiltered) condition. In both BB and HP condition, we found that SRTs for yNH were significantly lower than for oNH, and that SRTs for oNH were significantly lower than oHI (although the latter was not robust for the HP condition when outlier subjects were excluded). In contrast, in the LP condition, there was no significant difference between groups. These results can be best understood by considering the PTAs of the participants, which differed mainly in the high frequencies between the three groups (Fig. 2), therefore primarily limiting their processing capacities in that region. To further understand the engaged processes, we examined the relationships between SRTs across the different frequency filtering conditions. These correlation analyses suggest that SPiN performance of oNH and oHI individuals would be degraded by different components of hearing. The correlation between SRTs in BB and HP conditions observed for oHI individuals suggests that their deficits are primarily limited by differences in hearing thresholds in HFs. Consistent with that interpretation, we found a significant correlation between SRT in the HP condition and hearing thresholds in HFs for the oHI group, but not for the oNH group in which individuals among that group exhibit similar, close-to-normal thresholds (Fig. S1B, right panels). In contrast, the correlation between SRTs in BB and LP conditions observed for oNH, who have similar hearing thresholds at all frequencies, suggests that their deficits would be primarily linked to their processing of LFs, possibly related to TFS processing. In sum, the correlational analysis based on the three conditions portraits various deficit profiles for oNH and oHI listeners, but the results highlighted by the psychophysical and neural markers considered in that study provide important information to go beyond these observations (see below).

### FFR_env_ to RAM tone: a proxy of phase-locking capacities to ENV in the high frequencies

First, our results with the 4-kHz RAM tone replicate previous findings: FFR_env_ (often referred to as EFR in prior studies) led to a clear response indicating strong phase-locking to the rectangular envelope fluctuations of the sound (f0 of 110-Hz), with strong peaks related the first harmonics, from #1 to #4 (Vasilikov et al., 2021; Keshishzadeh et al., 2020; Mepani et al., 2021; Van der Biest et al., 2023). As in previous studies (Vasilikov et al., 2021; Keshishzadeh et al., 2020), we found clear and significant differences of FFR_env_ strength between all three groups of listeners (Fig. 4A), indicating effects of both age and hearing loss on ENV coding. Correlational analyses were partly consistent with these results. FFR_env_ was strongly negatively correlated to hearing thresholds (r=-0.59, *p* < .001), but the correlation with age was not significant (p > .05). The large variability in hearing thresholds among older participants likely hindered the relationship with age. Although CS is expected to increase with both age and hearing loss, the present findings are in line with previous studies showing that correlations between FFR_env_ and hearing thresholds can be observed, but not between FFR_env_ and age directly (Keshishzadeh et al., 2020). These results suggest that FFR_env_ is primarily sensitive to the impact of CS on ENV processing in the high frequencies (Vasilikov et al., 2021). The contribution of FFR_env_ to account for the variability in SRTs is further discussed with respect to the multiple regression analyses (see below). In contrast, and as expected since the 4-kHz carrier tone is at a frequency above the human limit of phase-locking to TFS (Verschooten et al., 2019), the FFR_tfs_ obtained with the RAM tone does not show phase-locking locking to the modulation frequency and its harmonics (Fig. 4A).

### FFR_tfs_ to a SM_harm_ stimulus: a proxy of phase-locking capacities to TFS in the low frequencies

Results obtained with the SM_harm_ stimulus show that the FFR was driven by both the ENV and the TFS components contained in that low-frequency stimulus. As for the RAM tone, significant differences observed between the three groups again indicate that FFR strength is negatively impacted by both hearing loss and age. The SM_harm_ also led to a FFR showing clear phase-locking to the envelope of the stimulus at 110 Hz, which corresponds to its fundamental frequency (FFR_env_; see Fig. 5A), and the first 3-4 harmonics, as observed with the RAM tone. However, the amplitude of that response was not as high as for the RAM-FFR_env_, a result that can be easily understood given that the temporal envelope of the SM_harm_ is not defined as strongly as for the RAM tone where by construction the envelope amplitude is decreased by 95% at every period. As with the RAM tone (see above), SM-FFR_env_ was correlated to hearing thresholds (r=-0.69, *p* < .001) but was not correlated to age (p > .05). Compared to the RAM stimulus that did not yield any FFR_tfs_, the SM_harm_ stimulus led to a measurable FFR_tfs_. It evidenced a clear harmonic structure, with the harmonic of highest amplitude matching harmonic #4 (440 Hz) corresponding to that nearest to the first formant peak (or here the ‘spectral modulation in the acoustical waveform. This observation is consistent with previous reports (e.g. Krishnan, 2002; Aiken & Picton, 2008; Ananthakrishnan et al., 2016, 2017).

We found that SM-FFR_tfs_ was correlated to hearing thresholds (r=-0.38, *p* = .009), but that it was also strongly correlated to age, a correlation that remained significant after adjusting for hearing thresholds (see Fig. 7A and 7B). This result reveals that SM-FFR_tfs_ is particularly sensitive to the impact of age, and that it more directly reflects TFS coding fidelity in the low frequencies without being additionally affected by other potential differences in OHC damage. These data provide additional evidence that aging impacts the strength of the TFS component of the FFR. Many previous works indeed reported effects of aging on FFR_tfs_ to tones or vowel-like stimuli. A strong magnitude reduction of FFRs with vowel-like stimuli (complex tones / vowel stimuli / voiced speech) was observed in older listeners with hearing loss compared to young normal-hearing listener, and that reduction was interpreted as primarily reflecting the effect of aging (Clinard and Cotter, 2015, Ananthakrishnan et al., 2016). A reduction in FFR_tfs_ strength was observed to static or dynamic pure tones (Clinard et al., 2010; Marmel et al., 2013; Märcher-Rørsted et al., 2022), as well as for more complex stimuli or speech-like sounds (Anderson et al., 2012, Bidelman et al., 2014; Mamo et al., 2016; Carcagno and Plack, 2020) for older compared to younger individuals, all having similar NH thresholds in the frequency range of the stimuli. FFRs to vowel sounds were also found to be smaller in HI compared to NH participants (Ananthakrishnan et al., 2016; Molis et al. 2023), and more specifically so concerning their TFS component. This evidenced reduction of FFR in older individuals is consistent with a degradation of TFS-coding with aging.

Yet, only few studies directly addressed the potential effect of CS on their results (Parthasarathy et al., 2020; Märcher-Rørsted et al., 2022). The analysis of our data in light of computational simulations suggest that the size of the difference in FFR_tfs_ strength between extreme individuals of ∼15-20 dB is consistent with a loss of ∼50-80% of HSR fibers, which is line with recent measurements obtained in histopathological human bone analyses (Wu et al., 2019, 2021). Importantly, computational simulations show that this decrease cannot be explained by OHC damage (Fig. 8), which would in contrast predict an increase – yet, minor – of the response. Even though CS since cannot practically be distinguished from IHC loss in the model used (Verhulst et al., 2018), recent studies already showed that a loss of IHC would have yielded much smaller effects (see Märcher-Rørsted et al., 2022). Moreover, another model of the auditory periphery (Zilany et al., 2014; URear) gave us decrease in FFR_tfs_ strength of comparable magnitude (not shown) to those reported here using the model from Verhulst et al. (2018). Therefore, the effects observed here are best accounted for by a decrease in the number of AN fibers, a prediction that appears to be robust.

The present model simulations suggest that a total removal of LSR and MSR fibers is not sufficient to predict any decrease in FFR strength of comparable magnitude to the one we observed empirically, and that a loss of ∼50-80% of HSR fibers seems required. This result appears consistent with previous reports that FFRs to high sound levels would primarily result from neural encoding in HSR AN fibers through off-frequency contributions (Encina Llamas et al., 2019; Carcagno and Plack, 2020; Temboury-Gutierrez et al., 2024). If this hypothesis is true, adding high-frequency noise to mask off-frequency contributions of SM_harm_ stimuli could yield a different pattern of result, in particular the observed negative correlation with age could be different (Carcagno and Plack, 2020). However, all of this remains to be addressed empirically.

### Integrating ENV-based and TFS-based markers: age-related effects reflecting a common deficit in neural phase-locking

An important result that emerged from our analyses concerns the fact that the age-FFR_tfs_ relationship was specifically and robustly driven by the magnitude of the 4^th^ harmonic of the FFR_tfs_ or, alternatively, by the sum of the magnitude of harmonics #4-#7 (Fig. 7). In sharp contrast, there was no correlation between age and the magnitude of the lower harmonics #1-#3 either considered individually or as a sum. This result suggests that age, and thus CS, would primarily impact phase locking capacities of cochlear neurons or AN fibers at quite high rates of ∼500 Hz. This is in line with previous studies that reported a decrease in FFR_tfs_, in which the frequency static or dynamic tones was above 500 Hz (Parthasarathy et al., 2020; Märcher-Rørsted et al., 2022). Consistent with this result, it is interesting to note that the correlation between RAM-FFR_env_ and SPiN scores reported in Mepani et al. (2021) was found to be specifically driven by the high harmonics (harmonics #3-#5) of the FFR_env_, corresponding to frequencies between 360 and 600 Hz. High-frequency carrier sounds modulated in amplitude and complex tones with low frequency components both monitor the capacity of auditory neurons to phase-lock to fast temporal information, the only difference being that this tracking concerns the ENV component of sound for neurons tuned to high CFs vs. the TFS component of sound for neurons tuned to low CFs, respectively, If this reasoning is correct, we would expect a relationship between the strength of the RAM-FFR_env_ and the strength of the SM-FFR_tfs_. This prediction was strongly supported by our data: we found a strong correlation between the two markers (p< .01), and more strikingly, this correlation was driven by the summed magnitude of the high harmonics (#4-#7) in both markers (r=0.53, *p* < .001), and even more precisely by the magnitude of harmonic #4 (r=0.44, *p* = .003). Even though this harmonic corresponds to the one of highest amplitude in the SM stimulus, making it difficult to disambiguate the specific role of its frequency from the fact that this is also the one mostly contributing to the input signal, it is important to note that the harmonic #4 in the RAM tone has no such dominating role in the input signal. Therefore, we believe that these correlations may rather suggest that the impact of age in low and high frequencies reflects a common decline in neural phase-locking for this frequency of ∼500 Hz. Beyond this finding, these results also nicely illustrate that although TFS and ENV coding are considered as separate dimensions before and within the cochlea, they are then merged into a single firing rate code. From a more practical perspective, our results suggest that considering FFR to complex low-frequency tones would constitute an efficient strategy to characterize slow as well as fast neural phase-locking capacities (Chauvette et al., 2022). Indeed, previous studies reported that it was not possible to obtain reliable FFRs using RAM tones or noises made of low-frequency carriers, suggesting that it might be best to limit RAM-FFR_env_ markers for the monitoring of CS to high-frequency carriers (Keshishzadeh et al., 2020).

We more globally characterized the relationships between the FFRs obtained with the RAM tone and the SM_harm_, as these stimuli are designed to differently engage processing of ENV / TFS cues in distinct frequency regions. There was a strong and significant correlation between the strength of the SM-FFR_env_ and the strength of the RAM-FFR_env_ (Fig. S3), suggesting that the deficit in ENV processing between the low and high frequencies would be strongly related. Since they rely on carriers that recruit specifically low or high frequencies, these responses likely reflect neural coding of sound engaged at different CFs. This result therefore suggests a strong relationship between the phase-locking capacities to temporal envelopes at different CFs on the cochlea. There was also also a strong correlation between SM-FFR_env_ and SM-FFR_tfs_, which primarily reflect coding fidelity of ENV or TFS cues of that stimulus, respectively (Aiken and Picton, 2008). This result thus suggests a link between the degree of deficit in ENV and TFS processing in the low frequencies, although more work is needed to really tease apart the relationship between these two metrics, which likely not independently reflect the coding of ENV and TFS features of the stimulus. However, these correlations, and in particular the one between RAM-FFR_env_ strength and SM-FFR_tfs_ strength were limited, also suggesting that they might reflect partially distinct aspects of auditory processing. This observation would be in line with Mai et al. (2023) who found FFR significantly decreased with age and high-frequency (≥ 2 kHz) hearing loss and also suggested distinct roles of FFR_env_ and FFR_tfs_.

Finally, our results suggest that the ability of neurons to extract fast rates of information in that neural code would be impaired by age-driven CS. Recent studies provide compelling evidence regarding for subcortical origins of speech-like FFRs and their relationship to SPiN capacities (Bidelman et al., 2020a, 2020b), and more specifically that FFRs to these range of high frequencies primarily result from neural generators at the level of the auditory nerve, compared to lower frequencies that would result from neural generators in the inferior colliculus and cochlear nucleus (Tichko and Skoe, 2017; Bidelman et al. 2018). The present results are in full agreement with this view. Since the computational models used to simulate FFRs do not include multiple generators, removing AN fibers impacted FFRs similarly at all frequencies, resulting in a similar decrease predicted for the TFS and the ENV components of the FFR (see Fig. 8), but we believe that this relationship is stronger in the case of TFS coding assessed in the low frequencies, where measurements are less additionally corrupted by the differences in the amount of outer-hair cell damage, since individuals have more comparable hearing thresholds in that range.

### Psychophysical detection of SM stimuli may not directly reflect TFS coding

Psychophysical detection thresholds for SM_harm_ stimuli were surprisingly highly variable across individuals, and led to non-significant differences across groups. Also, thresholds were higher (poorer) than in previous studies that used very similar conditions and stimuli, such as those of Bernstein et al. (2016), Miller et al. (2018). We verified the robustness of our results to different analysis methods (reconstruction of the psychometric functions vs. averaging of the last reversals in the adaptive track). First of all, we observed an overall correlation between SM_harm_ and SM_noise_ thresholds, suggesting a similarity between the thresholds measured with a SM envelope imposed on a noise vs. a harmonic carrier, which is consistent with the findings from Zaar et al. (2023a). Although previous studies observed higher (better) thresholds in the STM compared to the SM condition, consistent with the view that STM sounds provide an additional temporal cue for detecting the modulation compared to SM sounds that only carry a spectral, TFS-based cue (Miller et al., 2018; Zaar et al., 2023a), here we found that the thresholds with the STM_harm_ stimulus were higher and more variable, especially for yNH individuals, and did not correlate with thresholds for SM_harm_. It could be that our STM_harm_ stimuli, which included sidebands of noise, sounded very different from SM_harm_ and SM_noise_ to the participants and could therefore introduce some confusion regarding the task they had to perform. Overall, our empirical measurements show that the correspondence between the individual thresholds across the different stimuli conditions only lead to moderate correlation coefficients, which makes the thresholds obtained for the three types of stimuli not directly comparable. Thus, we mostly considered in our analyses the thresholds obtained with the SM_harm_, as this is the stimulus used for the FFR measurements, and because they correlated with thresholds obtained for SM_noise_, which was generally the stimulus used in previous works showing correlations with SRTs (Miller et al., 2018).

Contrary to previous works showing a correlation between SM thresholds and SRTs, we did not find any significant relationship between the two (Bernstein et al., 2013; 2016; Miller et al., 2018). We believe that there are a number of experimental parameters that distinguish our study from previous studies, which may explain this apparent discrepancy. First of all, these results might be very sensitive to the type of SPiN task. Zaar et al. (2023a) found correlations between STM thresholds and SRTs when assessed in a complex multi-talker spatial SPiN task (speech and multi-talker babble with masker speakers located at different spatial positions), but not in a co-located SPiN task (speech and speech-shape noise colocated in space) that was comparable to previous works (Miller et al., 2018). Several other experimental aspects may also play a role: the age and degree of hearing loss variability of the cohort of participants included, as well as the sound level at which they were tested. Here, SRTs and S(T)M detection thresholds were measured with stimuli at the same intensity for all participants, while in previous studies there was a level compensation based on the audiogram of the participant. Bernstein et al. (2016) and Miller et al. (2018) measured SPIN with hearing aids and Bernstein et al. (2013) compensated for hearing loss by presenting the sentences at higher intensity (above 90 dB HL). Zaar et al. (2023a) also tested HI participants with some compensation of their hearing loss. With our procedure, some frequency portions of the stimuli could not be processed in listeners, and the extreme case this resulted in an overall level that was not loud enough for a few participants with highest thresholds. Furthermore, previous studies such as Miller et al. (2018), only included HI listeners with large differences in their degrees of hearing loss and covering a wide range of ages. The present study included participants with NH and the cohort of HI listeners had only moderate hearing loss and a quite narrow age range. All these differences could be responsible for the differences between our observations and previous results.

### Results highlight the contribution of FFR_env_ for better predicting speech-in-noise intelligibility but fail to reveal further benefit of considering FFR_tfs_

We investigated the extent to which the neural and psychophysical measures provided by these experiments could predict the variability in SRTs assessed in the same listeners. First, analyses highlighted robust negative correlations between RAM-FFR_env_ and SRT in the BB as well as in the HP condition, replicating previous studies. However, there was no robust correlation between SM-FFR_tfs_ and SRT, specifically in the LP condition that was designed to maximize this potential relationship. Recent studies suggest that it is not easy to capture that the contribution of TFS in SPiN intelligibility. SPiN processing, even implemented in a laboratory-based task, necessarily engages a range of bottom-up and top-down cognitive processes that extend far beyond purely sensory mechanisms, and only a limited amount variance could be accounted for by any ideal marker of CS. Many recent studies suggest that cognitive factors could even play a larger role in accounting for the variability in most SPiN tasks, as compared to CS (Parthasaraty et al., 2020; Camerer et al., 2019; Gómez-Álvarez et al., 2023; Zink et al., 2024). Moreover, although our SPiN test used an LP filtering condition to favor the processing of TFS cues, this test might not be the most appropriate. Another aspect already discussed in the literature is the type of noise used for the SPIN test. Several studies showed that the role played by TFS cues is more easily measurable in the context of speech masked by multiple speakers, as compared to the case of a single speaker masked by signals with a complex, non-periodic TFS structure such as white noise or speech-shaped noise (Bharadwaj et al., 2019; Ruggles et al., 2012; Zeng et al., 2005). Also, the use of speech digits instead of the sentences of the Matrix test, could result in less recruitment of cognitive aspects that could impact our results (Parthasarathy et al., 2020; Ruggles et al., 2012). Won et al. (2016) successfully evidenced a direct relationship between the fidelity of formant encoding in FFR_tfs_ (estimated by taking the difference between the estimated peak on each subject and theoretical formant peak using PRAAT) and vowel confusion matrices measured in the same individuals; unfortunately, the same study did not test the extent to which it would contribute to SPiN variability in a more complex task. In sum, considering SPiN tasks with very simple speech stimuli maximizing the importance of TFS cues, such as isolated vowels or concurrent vowel trajectories (Woods & McDermott, 2015), would help highlighting the extent to which it may be accounted for by FFR_tfs_.

### Further research is required to use psychophysical SM detection thresholds as a direct reliable behavioral proxy of TFS coding fidelity

One possible explanation for the absence of a relationship between SM psychophysical thresholds and FFR_tfs_ is that these two measures may reflect different aspects of auditory processing. In our study, the SM-FFR was assessed using spectrally modulated sounds with 50% and 100% modulation depths to better capture formant encoding. We hypothesized that the amplitude differences between harmonics in the SM-FFR spectrum for these two modulation depths would be a better predictor of SM thresholds than absolute amplitude alone (cf. Won et al., 2016). However, no significant difference was observed in the FFR responses between the two modulation depths, leading us to use the average response as the neural metric. These results raise questions about whether psychophysical SM detection thresholds are sufficiently sensitive to reflect TFS processing. To address this, future studies might explore other psychophysical tests using stimuli that are more directly related to TFS processing, such as the TFS1 test (Sęk & Moore, 2012) or frequency modulation (FM) thresholds using low-frequency carriers (Borjigin et al., 2022; Parthasarathy et al., 2020). Moreover, psychophysical experiments aimed at characterizing sensory capacities should account for potential cognitive and attentional factors that could influence performance. Borjigin et al. (2022) demonstrated how attentional lapses during psychophysical tasks can account for nearly 50% of the variance in amplitude modulation (AM) threshold measurements, underscoring the importance of controlling for these factors in future studies. Despite the lack of correlation between the FFR and psychophysical measures in our study, this does not undermine the possibility that either psychophysical measures of SM detection/discrimination or FFRs to SM stimuli could be viable tools for assessing TFS coding and/or CS in the future, depending on the research context. For instance, Märcher-Rørsted (2022) provided strong evidence that FFRs to pure tones are reduced with age, suggesting that simple tones could be used to monitor CS. However, the potential for using complex SM stimuli in psychophysical tests warrants further investigation. These stimuli are particularly appealing because they serve as parameterized models of vowel sounds (Ponsot et al., 2021), which could improve the link between TFS coding and performance on SPiN tasks.

## Acknowledgments

We acknowledge the contribution of Heleen Van Der Biest and Saartje Vanaudenaerde for their effort in collecting the data of this paper. We would also like to thank Fotis Drakopoulos and Sarineh Keshishzadeh for their advices and help when designing and conducting the first EEG experiments using STM sounds. This work was supported by a postdoctoral fellowship by the Fondation pour l’Audition [FPA 2020-005F2], the European Research Council [ERC-StG-678120 to S.V., RobSpear] and the ANR [ANR-22-CE28-0010 to E.P., InspectSyn].

## Data availability statement

The data generated and analyzed during this study will be made publicly available on the Open Science Framework (OSF) upon publication.

## Supplementary material

**Figure S1.**
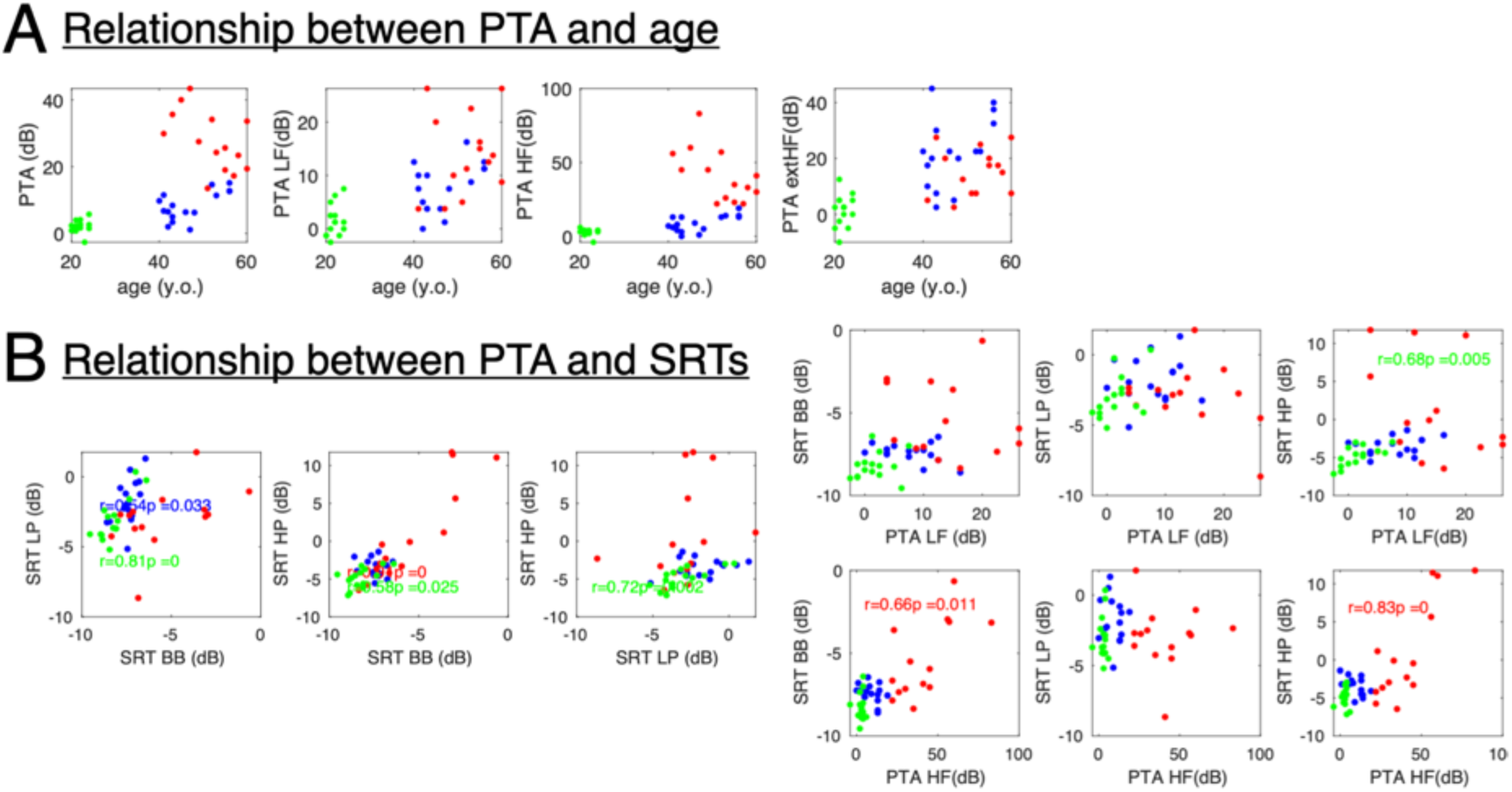
(A) Relationships between age and PTA computed using all frequencies (0.125-8 kHz; PTA), low frequencies (0.125-1 kHz; PTA LF), or high frequencies (2-8 kHz; PTA HF). (B) Left panels show the relationships between SRTs across the different frequency filtering conditions (BB, HP and LP; see the main text for details) and right panels between SRTs and PTA computed using different frequency ranges; significant correlations (pearson, uncorrected) are shown in the corresponding panels, with colors corresponding to each specific group

**Figure S2.**
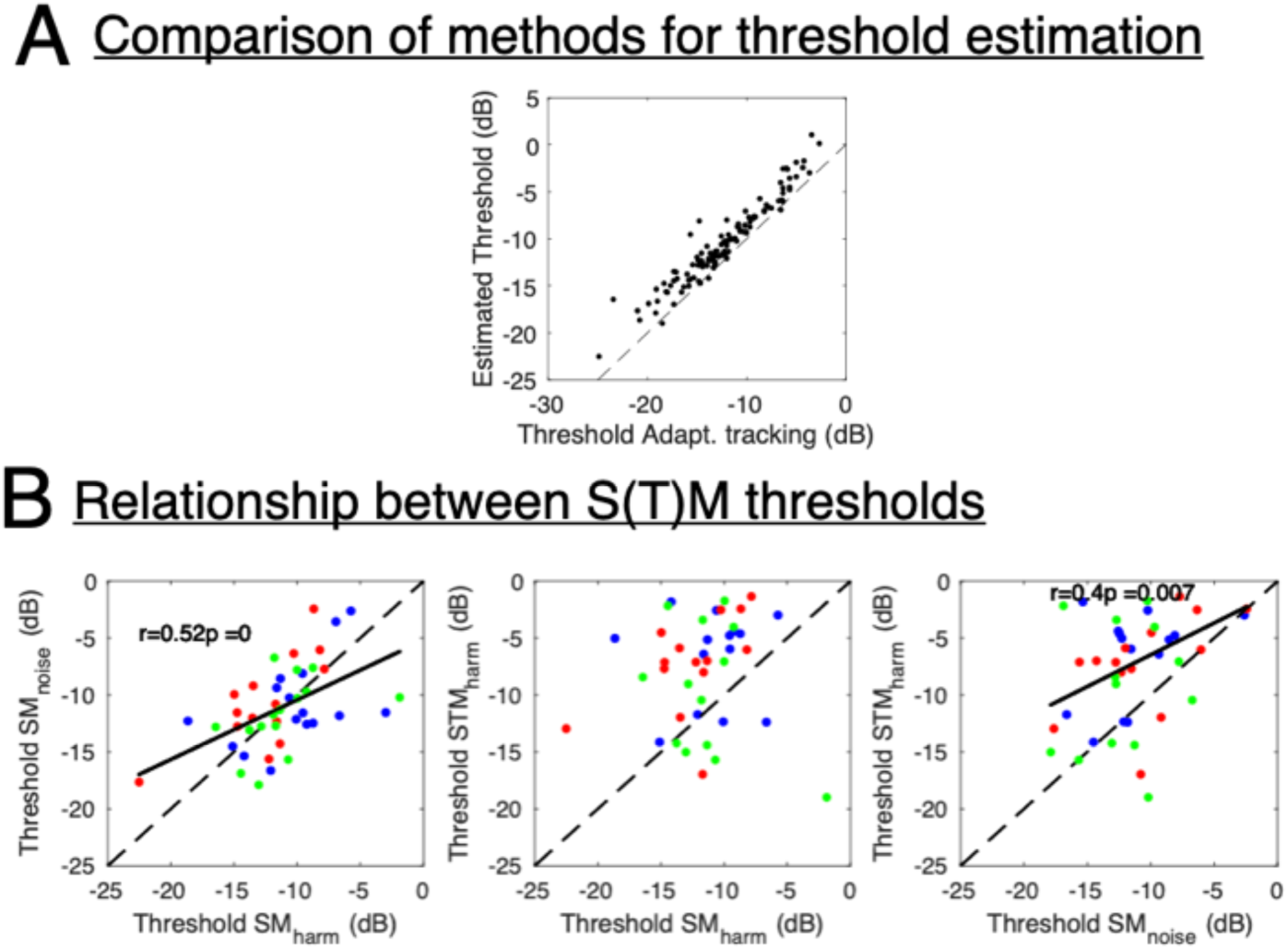
(A) Comparison of thresholds obtained from psychometric function reconstruction (the preferred method for the present study) and simple averaging of the last six reversals of the adaptive tracks, computed for all individuals and conditions in which both methods could be used alternatively. A strong correlation was obtained (p < 0.001) between both types of estimation, and as expected (a higher point is tracked on the psychometric function), the thresholds estimated using the psychometric function reconstruction method were higher than those obtained from averaging over the final 6 reversals (p < 0.001). (B) Relationship across thresholds derived for the three types of stimuli. Significant correlations (pearson, uncorrected) are shown in the corresponding panels (all individuals aggregated).

**Figure S3.**
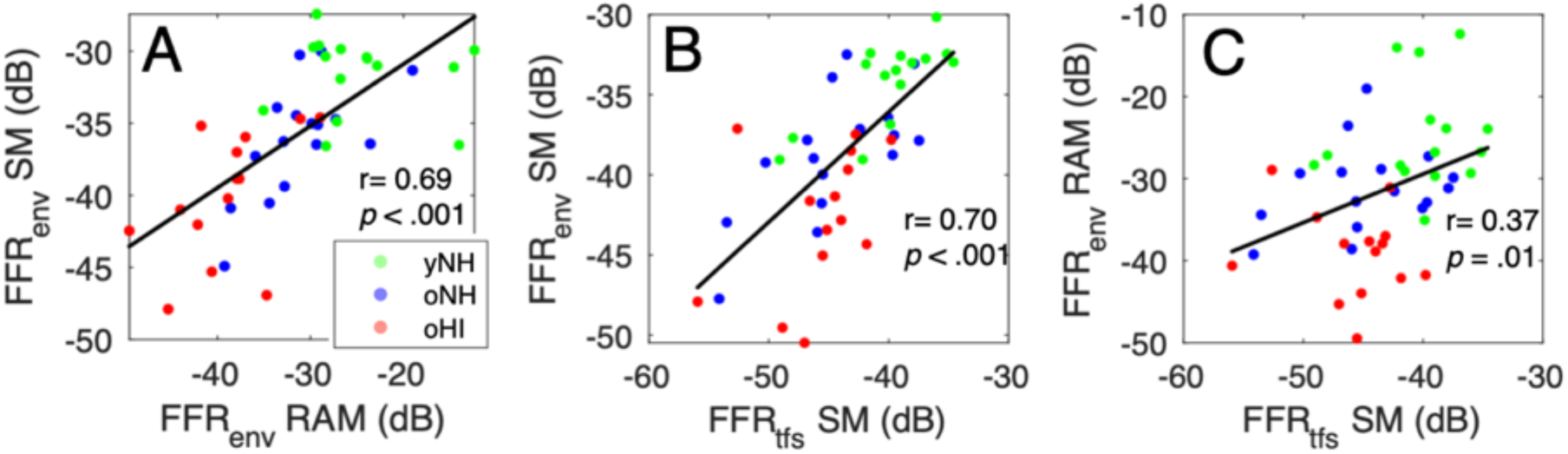
Relationships between SM-FFRenv and RAM-FFRenv (A, r=0.69, p<.001, all individuals; in black, the linear regression line), between SM-FFRenv and SM-FFRtfs (B, r=0.70, p<.001), and between RAM-FFRenv and SM-FFRtfs (C, r=0.37, p=0.01).

**Figure S4.**
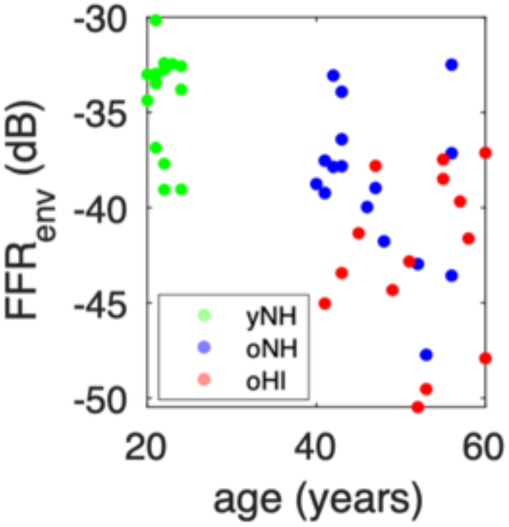
No correlation was observed between FFRenv to the RAM tone and age when considering oHI and oNH individuals together (p > .05).

**Table S1.**
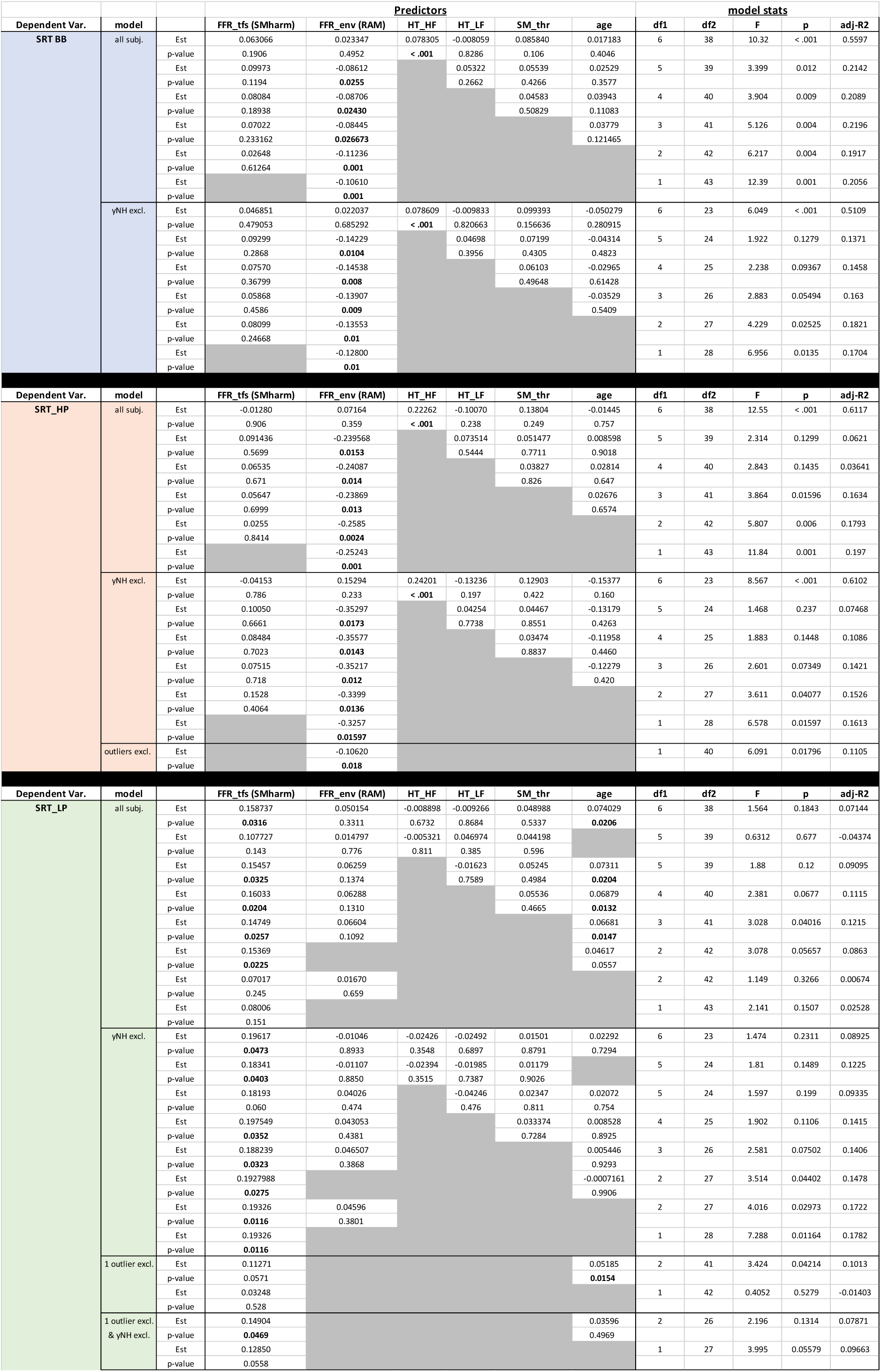
Results of multiple linear regression models conducted to assess the contribution of the different predictors derived in the present study.

